# esBAF and INO80C fine-tune subcompartments and differentially regulate enhancer-promoter interactions

**DOI:** 10.1101/2025.09.16.676562

**Authors:** Braulio Bonilla, Benjamin J. Patty, Snehal V. Sambare, Job Dekker, Thomas G. Fazzio, Sarah J. Hainer

**Affiliations:** Department of Biological Sciences, University of Pittsburgh, Pittsburgh, PA, 15260 USA; Department of Systems Biology, University of Massachusetts Chan Medical School, Worcester, MA, 01655, USA; Howard Hughes Medical Institute, Chevy Chase, MD, 20815, USA; Department of Molecular, Cell, and Cancer Biology, University of Massachusetts Chan Medical School, Worcester, MA, 01655, USA; UPMC Hillman Cancer Center, Pittsburgh, PA, 15232, USA

**Keywords:** remodelers, chromatin, subcompartments, enhancer-promoter interactions, stem cells

## Abstract

The genome is compacted in the nucleus through a hierarchical chromatin organization, ranging from chromosome territories to compartments, topologically associating domains (TADs), and individual nucleosomes. Nucleosome remodeling complexes hydrolyze ATP to translocate DNA and thereby mobilize histone proteins. While nucleosome remodeling complexes have been extensively studied for their roles in regulating nucleosome positioning and accessibility, their contributions to higher-order chromatin architecture remain less well understood. Here, we investigate the roles of two key nucleosome remodelers, esBAF and INO80C, in shaping 3D genome organization in mouse embryonic stem cells. Using Hi-C, we find that loss of either remodeler has minimal effects on global compartment or TAD structures. In contrast, subcompartment organization is notably altered, suggesting that esBAF and INO80C contribute to finer-scale chromatin topology. To overcome the limited resolution of Hi-C for detecting regulatory loops, we employed promoter capture Micro-C (PCMC), which revealed that the loss of esBAF or INO80C alters a subset of promoter anchored looping interactions. Although these changes occur at distinct genomic loci for each remodeler, the affected sites are commonly enriched for bivalent chromatin regions bound by OCT4, SOX2, and NANOG (OSN), as well as BRG1 and INO80 themselves. Together, our findings reveal that esBAF and INO80C selectively influence subcompartment identity and enhancer–promoter communication at key regulatory loci, highlighting a previously underappreciated role for nucleosome remodelers in higher-order chromatin organization.

## Introduction

Chromatin is organized across multiple length scales, from individual nucleosomes, the fundamental repeating unit, to higher-order domains and long-range contacts that underlie nuclear architecture (Dekker & Mirny, 2024; Dekker et al., 2013; Fransz & de Jong, 2011; Lieberman-Aiden et al., 2009; Luger et al., 1997; Schoenfelder & Fraser, 2019). Local nucleosome positioning governs DNA accessibility and thereby exerts some control over regulatory element activity, including permitting access to transcription factors and RNA Polymerase II activity (4D Nucleome Consortium et al., 2024; Buenrostro et al., 2013; Dekker & Mirny, 2024; Dixon et al., 2015; Kornberg & Lorch, 1999; Kouzarides, 2007; Lieberman-Aiden et al., 2009; G. Li & Reinberg, 2011; Rowley et al., 2017; Sexton et al., 2016). Advances in high-resolution chromatin conformation assays (Hi-C, Micro-C), ligation-independent contact mapping, and super-resolution imaging have revealed a dynamic 3D genome at different length scales composed of compartments at the sub Mb to Mb scale, subcompartments at the sub Mb scale, topologically associating domains (TADs) at the scale of tens to hundreds of kb, and enhancer–promoter loops between cis-regulatory elements (Beagrie et al., 2017, 2023; Beliveau et al., 2015; Bintu et al., 2018; Fiorillo et al., 2021; Lieberman-Aiden et al., 2009; Maslova & Krasikova, 2021; McCord et al., 2020; Quinodoz et al., 2018; S. Wang et al., 2016). At the largest scale, chromosomes occupy distinct territories, which are subdivided into active (A) and inactive (B) compartments that reflect transcriptional activity and chromatin state (Lieberman-Aiden et al., 2009). In many cell types, these compartments reflect a spatial segregation of heterochromatic domains, such as lamina-associated domains (LADs) and nucleolus-associated domains (NADs), from more transcriptionally active euchromatin. Finer-scale structures known as subcompartments have been more recently described, revealing additional layers of regulatory specificity (Rao et al., 2014). These subcompartments are defined by characteristic patterns of histone modifications, DNA accessibility, and spatial nuclear localization (Rao et al., 2014). Topologically associating domains (TADs) constrain chromatin contacts into self-interacting regions, limiting enhancer–promoter communication within their boundaries (Dixon et al., 2012; Nora et al., 2012). Chromatin loops further organize the genome by connecting distal enhancers and promoters, thereby shaping gene expression programs. Together, these structural layers provide a multi-scale framework for genome regulation. How local nucleosome remodeling events scale up to influence these higher-order folding patterns, however, remains poorly defined.

ATP-dependent nucleosome remodeling complexes, including SWI/SNF (esBAF in embryonic stem cells), ISWI, and INO80 families, are primary effectors of local chromatin structure: they reposition or evict nucleosomes, or alter nucleosome composition to modulate accessibility for transcription, replication, and repair (Clapier et al., 2017; Narlikar et al., 2013). In pluripotent cells, esBAF and INO80C support open chromatin at enhancers and precise positioning of promoter-proximal nucleosomes, activities that promote enhancer–promoter communication and transcriptional robustness (Ho et al., 2011; Hainer & Fazzio, 2015; Krietenstein et al., 2016; Wang et al., 2014). Nucleosome depleted regions, observed at cis-regulatory elements, are generated by nucleosome remodelers. Recently, these nucleosome depleted regions have been shown to be sufficient to make a domain boundary *in vitro*, linking remodelers to higher order chromatin structure (Aljahani et al., 2024; Oberbeckmann et al., 2024). In addition, remodeler activity at insulator elements can influence nucleosome architecture in ways that impact binding of CTCF, a key regulator of higher-order chromosome folding (Barisic et al., 2019). Despite these well-established local roles and known connections with CTCF, the extent to which remodelers shape megabase-scale genome folding including compartments, subcompartments, TAD insulation, and looping, remains unclear.

Perturbation of architectural factors such as CTCF or cohesin causes pronounced changes in loop and TAD organization yet often yields surprisingly modest immediate effects on global gene expression, underscoring the robust and context dependent relationship between structure and function (Dixon et al., 2012; Nora et al., 2012) Nora et al., 2017; (Rao et al., 2014). Studies that directly interrogate nucleosome remodeling complex function in 3D chromosome organization have reported variable outcomes: for example, BRG1 (the catalytic subunit of BAF) was implicated in establishment or maintenance of TAD boundaries in some cell types (Barutcu et al., 2016), whereas depletion of the same nucleosome remodeling complexes produced minimal TAD changes in mouse embryonic stem cells (Barisic et al., 2019), suggesting cell-type specificity and scale-dependent effects. Importantly, remodelers typically act locally, often repositioning only a handful of nucleosomes at a given regulatory element. This raises a key mechanistic question: to what extent do local remodeling events propagate to alter enhancer-promoter looping, subcompartment identity, or other fine-scale folding features that govern gene regulation?

Addressing this question requires coupling precise, often rapid perturbations of remodeler activity with high-resolution chromatin conformation mapping and complementary readouts of nucleosome positioning and the chromatin-bound proteome. Here, we apply Hi-C and promoter-capture Micro-C (PCMC) following depletion of BRG1 (esBAF) or INO80C in mouse embryonic stem (ES) cells to probe how remodeler loss affects folding at multiple scales. Although global compartmentalization and canonical TAD structures are largely preserved, we detect reproducible, locus-specific changes in subcompartment organization and enhancer– promoter looping particularly at bivalent loci bound by OCT4, SOX2 and NANOG. While *Smarca4* KD mainly decreases the number of loops at bivalent loci, *Ino80* KD increases the number of loops at these loci. Given that they act on distinct bivalent loci, esBAF and INO80C exert distinct, fine-scale control over chromatin architecture: local repositioning of a small number of nucleosomes at key regulatory elements can measurably rewire higher-order contacts, with implications for transcriptional regulation in pluripotent cells.

## Methods

### Cell lines

E14TG2a (E14) mouse embryonic stem (ES) cells derived from male *Mus musculus* (RRID:CVCL9108; (Hooper et al., 1987)), were cultured under standard conditions. Cells were maintained on tissue culture-treated plates pre-coated with 0.2% gelatin and incubated at 37°C in a humidified atmosphere containing 5% CO₂. The culture medium consisted of high-glucose Dulbecco’s Modified Eagle Medium (DMEM; Sigma, D5796), supplemented with 10% fetal bovine serum (Sigma), 2 mM L-glutamine (Corning), 0.1 mM β-mercaptoethanol (Acros Organics), 1X nonessential amino acids (Corning), and leukemia inhibitory factor (LIF; prepared in-house). Cells were passaged every 48 hours using trypsin-EDTA (Corning, 25-053-Cl) at a dilution of approximately 1:8 in fresh medium. Mycoplasma contamination was routinely assessed by PCR, and culture space was treated with anti-mycoplasma solution (LookOut® DNA Erase Spray, Sigma) to minimize contamination risk. All experiments were conducted using ES cells between passages 18 and 26.

### esiRNA Knockdowns

Endoribonuclease-digested short interfering RNAs (esiRNAs) were generated as previously described (Calegari et al., 2002; Fazzio et al., 2008; Klein, Troy, et al., 2023; Yang et al., 2002). Briefly, target gene regions were selected based on sequence uniqueness using the DEQOR algorithm (Henschel et al., 2004). For esiRNA synthesis, T7 templates were generated using cDNA derived from wild-type murine E14 embryonic stem (ES) cells for *Smarca4* and *Ino80*, and from the pLJM1-EGFP plasmid (Addgene, Plasmid #19319) (Sancak et al., 2008), for control *EGFP* esiRNAs. The oligonucleotide sequences used to generate T7 templates are listed in Table S1. The dsRNAs were *in vitro* transcribed using T7 polymerase (homemade; 5.5 hr at 37°C, followed by an annealing stage lowering the temperature 0.1°C/s down to 25 °C), digested with ShortCut RNase III (NEB), then purified using a PureLink RNA Mini Kit (Invitrogen) with a modified purification strategy. Briefly, lysis buffer was added to a 1:1 volume ratio and isopropanol was then added to a final concentration of 40%. Following a 15 sec vortex, the samples were loaded to silica columns (Epoch Life Science) and centrifugated (1 min, 11,000 x*g*). Isopropanol was added to the flow-through at a 1:1 volume. Following a 15 sec vortex, the samples were applied to new silica columns and centrifugated (1 min, 11,000 x*g*). This was repeated until the entire sample was applied to the column. After a 500 µL wash with Wash buffer 2, the samples were eluted using ultrapure nuclease-free water (Ambion). Transient transfections were performed in 10 cm plates using Lipofectamine 3000 (Invitrogen). Briefly, 3.5 µg of esiRNAs and 25 µL of Lipofectamine were incubated in 2 mL of OptiMEM (Gibco) for 20 min. Then, 3.5X10^5^ cells in 4 mL of culture media were added to the transfection mixture and plated onto 10 cm plates. After 16-18 hr, the media was removed, and fresh media was added. Samples were harvested 48 hr post transfection using trypsin. Following a 1X PBS (Corning) wash, cells were processed or flash-frozen and held at −80°C.

### RT-qPCR

For *in situ* Hi-C experiments, validation of the KD efficiency was performed by RT-qPCR. Total RNA was extracted using TRIzol Reagent (Ambion) following manufacturer instructions. cDNA was generated using random hexamer primers (Promega, C118A) and 1 µg of RNA. qPCR was performed on a Roche Lightcycler 96 using SYBR green (KAPA), while *Gapdh* was used as a reference gene. Fold change was calculated using the ΔΔCT method (Livak & Schmittgen, 2001). The oligonucleotides used for RT-qPCR are listed in Table S1.

### Western blot

For the PCMC experiments, validation of KD efficiency was performed by western blot. Whole protein extraction from frozen cell pellets was performed using RIPA buffer (150 mM NaCl, 50 mM Tris-HCl pH 8.0, 1% NP-40, 0.5% sodium deoxycholate, 0.1% SDS) supplemented with freshly added 1 mM DTT and protease inhibitors (Thermo Scientific). 1X10^6^ cells were incubated with 70 µL of RIPA buffer for 20 min in ice, vortexing every 3 min. The lysates were cleared by centrifugation. 50 µg of protein were prepared with 4X Sample Buffer (200 mM Tris-HCl pH 6.8, 8% w/v SDS, 40% glycerol) and loaded onto 9% PAGE gels. The proteins were transferred overnight at 4°C and at 20V to a nitrocellulose membrane (BioTrace). Equal protein loading was assessed using REVERT 700 stain (LI-COR). After blocking for 1 hr at RT, the membranes were incubated with 1:1000 Polyclonal rabbit anti-INO80 (Invitrogen, PA5-65296) or 1:1000 monoclonal rabbit anti-BRG1 (Cell Signaling, D1Q7F) in PBS-T for 3 hr at RT. Secondary goat anti-rabbit antibody (LI-COR, 92632211; 1:10,000) was added in PBS-T for 1 hr at RT. Blots were analyzed using Odyssey M imager (LI-COR). Depletion efficiency was calculated using ImageJ (Schindelin et al., 2012).

### In situ Hi-C

*In situ* Hi-C was performed as described (Belaghzal et al., 2017). Briefly, 5X10^6^ cells were crosslinked using formaldehyde (1% final concentration) for 10 min at RT. The reaction was quenched using glycine (130 mM final concentration). After a PBS wash, the cell pellets were frozen and stored at −80°C. Cells were resuspended in ice-cold lysis buffer (10 mM Tris-HCl pH 8.0, 10 mM NaCl, 0.2 % NP-40) with protease inhibitors (Thermo Scientific) and incubated on ice for 15 min. The lysate was homogenized using dounce homogenization and nuclei were pelleted at 2,500 x*g* at RT. The nuclei were then washed with and resuspended in ice-cold NEBuffer 3.1 (NEB). SDS was then added (0.1% final concentration) and the nuclei were incubated at 65°C for 10 min. SDS was quenched by adding Triton X-100 (1% final concentration). *DpnII* (50,000 U, NEB) was added, and chromatin digestion was carried out overnight at 37°C. End repair was performed at 23°C for 5 hr using Klenow Fragment (3’→5’ exo-, 50 U, NEB), NEBuffer 3.1 (0.112 X), 28 nM of each dCTP, dGTP. dTTP, and biotin-14-dATP. Blunt ligation was performed at 16°C for 4 hr using T4 DNA ligase (60 U, Invitrogen) and ligation buffer (1X, Invitrogen), Triton X-100 (1%) and BSA (1 mg/mL). Crosslinks were reversed at 65°C overnight, and protein was degraded using proteinase K. The DNA was extracted using phenol-chloroform (Invitrogen). Biotin removal from unligated ends was performed at 20°C for 4 hr using 5 µg of DNA, 5 U T4 DNA Pol, NEBuffer 2.1 (1X, NEB), 25 nM dATP, and 25 nM dGTP. DNA was sheared to obtain 200-300 bp fragments sizes using a sonicator (Covaris).

DNA size fractionation was performed using AMPure XP magnetic beads (Beckman Coulter) as described by the manufacturer. End repair was performed at 20°C for 30 min using 50 µL of size-selected DNA elute, T4 DNA pol (7.5 U, NEB), T4 polynucleotide kinase (25 U, NEB), Klenow DNA pol (2.5 U, NEB), ligation buffer (1X, NEB). Biotin DNA pulldown was performed using 1 µg of end-repaired DNA, MyOne streptavidin C1-coated beads (Invitrogen), according to manufacturer instructions. The beads were resuspended in 41 µL TE. A-tailing was then performed at 37°C for 30 min, followed by addition of Klenow fragment (3’ -> 5’ exo-, 15 U, NEB), dATP (0.2 nM), and NEBuffer 2.1 (1X). After a wash with ligation buffer, the beads were resuspended in 36.25 µL ligation buffer. Adapter ligation was then performed at RT for 2 hr with the addition of T4 ligase (4 U), ATP (0.2 mM), paired ended adapters (5 µL of 15 µM annealed adapters, 5’ ligation buffer (2.75 µL). After washing twice with tween wash buffer (TWB; 5 mM Tris-HCl [pH=8.0], 0.5 mM EDTA, 1 M NaCl, 0.05% Tween), the beads were resuspended in Binding Buffer (10 mM Tris-HCl pH 8.0, 1 mM EDTA, 2 M NaCl). Following a wash with NEBuffer 2.1, beads were resuspended in 20 µL of NEBuffer 2.1. The library of ligated fragments was then PCR amplified for 6 cycles using 6 µL of beads, 2 µL of 10 µM adapter primers (G19-PE Primer 1.0 and 2.0), 25mM dNTPs, 10X PfU buffer, and 2 µL PfuUltra in a 30 µL reaction and the following conditions: 98°C for 3 min, 5 cycles of: 98°C for 20 sec, 60°C for 30 sec, and 72°C; and a final amplification at 72°C for 5 min. Finally, the PCR products were combined, and primer dimers were removed using AMPure XP beads. The libraries were pooled and sequenced to ∼1 billion uniquely mapped reads using Illumina NextSeq500.

### Hi-C data analysis

Raw fastq files for each replicate were aligned to the mm10 genome using BWA (H. Li & Durbin, 2009, 2010) and the resulting sam files were downsampled to the smallest for each replicate using SAMtools (Danecek et al., 2021). The datasets were then processed with different tools to obtain matrices in the desired formats. Juicer 2.0 (Durand et al., 2016) was used to obtain matrices in the Juicer format (hic files). Matrices from the two biological replicates were combined using the mega.sh script from Juicer. nf-core/hic (Servant et al., 2023) was used to obtain matrices in the cooler format (cool files) (Abdennur & Mirny, 2020) Replicas were combined using the merge command from the cooler library (Abdennur & Mirny, 2020). HOMER (Heinz et al., 2010) was used to generate tag directories, the default HiC pipeline and options from HOMER were used to perform downstream analyses. Tag directories from two biological replicas were combined using the makeTagDirectory command from HOMER. To obtain HiCExplorer (Ramírez et al., 2018) format matrices (h5 files), we converted raw Juicer matrices of the corresponding resolutions to FAN-C (Kruse et al., 2020) format using the FAN-C command fanc to-fanc, the FAN-C matrices were then exported to cooler matrices using fanc to-cooler and later converted to HiCExplorer format using the HiCExplorer command hicConvertFormat. Finally, we normalized the matrices using the hicCorrectMatrix command from HiCExplorer. All genomic feature enrichment analyses and genomic feature list manipulations were performed using bedtools (Quinlan & Hall, 2010).

#### Compartment analysis

To identify compartments, we first performed a PCA analysis with HOMER using the runHiCpca.pl script (default settings) on the combined tag directories at a 25 kb resolution, then the compartments were called using findHiCCompartments.pl (default settings). Pearson correlation matrices were obtained using 250 kb resolution FAN-C matrices and the fanc compartments command. Browser tracks were generated in UCSC genome browser.

#### Subcompartment analysis

Subcompartment analysis was performed using CALDER2 (Liu et al., 2021) at 50 kb resolution on matrices generated using Juicer2. To compute the subcompartment change percentages, we recorded the subcompartment assignment of the central position of each subcompartment found in *Smarca4* KD or *Ino80* KD samples and compared the assignment in the control sample (**Fig. 2A**). To analyze the ratio of subcompartment switches associated with specific genomic features we, similarly, recorded the subcompartment assignment for each location associated with each feature in *Smarca4* KD or *Ino80* KD samples and compared it to the one observed in the control sample (**Fig. 2B; S3A**). To obtain the observed/expected ratio of chromatin opening or closing, for each feature, we divided the number of subcompartments that switched to a more active state or a more inactive state over an average of 1,000 randomized genomic locations (**Fig. 2C-G; S3D,F-H**). Randomized genomic locations were generated using bedtools shuffle (Quinlan & Hall, 2010) for each genomic feature analyzed. Browser tracks were generated in UCSC genome browser.

#### TAD analysis

TAD boundaries were calculated with the hicFindTADs command from HiCExplorer v3.7.2, using 10 kb resolution HiCExplorer matrices and the following settings: correctForMultipleTesting fdr, minBoundaryDistance 80000, minDepth 80000, maxDepth 800000, step 25000, and thresholdComparisons 0.05. Differential TADs were calculated using two independent approaches: 1) with the hicDifferentialTAD command from HiCExplorer v3.7.2 using the results from the aforementioned hicFindTADs analysis and the following settings: mode all, modeReject all, pValue 0.05; 2) TADCompare (Cresswell & Dozmorov, 2020) with default settings on Juicer-generated 25 kb resolution raw matrices. Insulation was calculated on 25 kb resolution cooler format matrices using the _find_insulating_boundaries_dense command from cooltools v0.7.0 at 125 kb resolution. To compare the insulation values at TAD boundaries associated with BRG1 or INO80 binding, we used a list of TAD boundaries obtained with hicFindTADs and intersected them with CTCF, BRG1 or INO80 binding locations using bedtools.

#### Loop analysis

Hi-C loops were identified using Mustache (Roayaei Ardakany et al., 2020) using 10 kb resolution, FDR=0.01, and otherwise default settings. Differential Hi-C loops were obtained using diff_mustache.py (Roayaei Ardakany et al., 2020) using 10 kb resolution and an FDR=0.01.

### Analysis of public ChIP-seq data

ChIP-seq datasets were obtained from the NCBI Sequence Read Archive (SRA; Table S2). Analysis of these data sets was performed as follows: fastq files were trimmed using trimmomatic 0.38 (options: -phred33 ILLUMINACLIP:TruSeq3-PE.fa:2:30:10 LEADING:3 TRAILING:3 SLIDINGWINDOW:4:15 MINLEN:36). The trimmed files were then mapped to the mouse genome (mm10) using bowtie (options: -n 1 --best -l 75 --strata -m 1; the flag -- chunkmbs 2000 was added for paired-end fastq files) (Langmead et al., 2009). Uniquely mapped reads were then filtered for MAPQ ≥ 10 using SAMtools (H. Li & Durbin, 2009). HOMER (Heinz et al., 2010) was then used to call peaks (findPeaks command, default options).

### CUT&RUN

CUT&RUN was carried out as previously described (Hainer & Fazzio, 2019; Hainer et al., 2019; Klein, Lardo, et al., 2023; Lardo & Hainer, 2022; Patty & Hainer, 2021; Skene & Henikoff, 2017). Briefly, nuclear extraction was performed by incubating 100,000 cells with a hypotonic lysis buffer (20 mM HEPES–KOH, pH 7.9, 10 mM KCl, 0.5 mM spermidine, 0.1% Triton X-100, 20% glycerol, and protease inhibitors). The nuclei were then bound to lectin-coated concanavalin A magnetic beads (40 µL bead slurry per 100,000 nuclei, Polysciences), treated with blocking buffer (20 mM HEPES, pH 7.5, 150 mM NaCl, 0.5 mM spermidine, 0.1% BSA, 2 mM EDTA, and protease inhibitors), and washed with washing buffer (20 mM HEPES, pH 7.5, 150 mM NaCl, 0.5 mM spermidine, 0.1% BSA, and protease inhibitors). Primary antibody (BRG1: Bethyl, A300-813A lot 5) incubation was performed in wash buffer for 1 hr at room temperature. The nuclei were then incubated with Protein A/Protein G-MNase (pA/G-MN, homemade) in wash buffer for 30 min at room temperature. The reactions were chilled to 0°C on an ice waterbath followed by the addition of CaCl_2_ (3 mM) to activate MNase digestion. The reaction was quenched with EDTA (20 mM) and EGTA (4 mM) after 15 min. MNase-treated genomic DNA from *S. cerevisiae* (1.5 pg) was added as spike-in control. DNA was released using RNaseA treatment and centrifugation. Libraries for sequencing were built by subsequently performing end repair and adenylation, NEBNext stem-loop adapter ligation and AMPure XP bead (Beckman Coulter) purification. The libraries were then PCR amplified by 14 cycles, and the products were purified with AMPure XP. Finally, the libraries were pooled and sequenced to at least ∼10 million uniquely mapped reads using Illumina NextSeq500.

### CUT&RUN data analysis

CUT&RUN data was processed and analyzed as described previously (Klein, Lardo, et al., 2023; Patty & Hainer, 2021). Raw fastq files were trimmed to 25 bp and mapped to the mouse genome (mm10) using bowtie2/2.4.1 (options -q -N 1 -X 1000) (Langmead & Salzberg, 2012). Duplicate reads were removed using Picard/2.18.12 (Picard toolkit, 2019) while reads with low mapping quality were removed using SAMtools (MAPQ ≥ 10) (H. Li et al., 2009). Fragment sizes < 120 bp corresponding to BRG1 binding were then obtained using SAMtools and a custom awk script. HOMER (Heinz et al., 2010) was used to call peaks (findPeaks command, default options).

### Promoter capture Micro-C (PCMC)

The Micro-C library was performed using the Dovetail® Micro-C Kit according to the manufacturer’s protocol. Briefly, chromatin was fixed with disuccinimidyl glutarate (DSG) and formaldehyde. The crosslinked chromatin was then digested *in situ* with micrococcal nuclease (MNase). Following digestion, the cells were lysed with SDS to extract the chromatin fragments and the chromatin fragments were bound to chromatin capture beads. Next, the chromatin ends were repaired and ligated to a biotinylated bridge adapter followed by proximity ligation of adapter-containing ends. After proximity ligation, the crosslinks were reversed, the associated proteins were degraded, and the DNA was purified used as input for sequencing library preparation using Illumina-adapters. Biotin-containing fragments were isolated using streptavidin beads prior to PCR amplification. The library was then enriched for promoter regions using Dovetail® Pan Promoter Enrichment Kit (CP3-PP-002) following manufacturer instructions. The libraries were pooled and sequenced using Illumina NextSeq500.

### Promoter capture Micro-C (PCMC) analysis

Promoter-capture micro-C matrices were downsampled across replicas to obtain matrices with equal contacts. We obtained loops using CHiCAGO (Cairns et al., 2016) pipeline with 20 kb resolution. The loops from both replicas were combined using pgltools intersect. These loops were compared to the Hi-C loops obtained from the control samples using bedtools (Quinlan & Hall, 2010). Differential loop analysis was performed using the CHiCAGO (Cairns et al., 2016) pipeline on combined replicas. We considered p < 0.5 as the threshold for loop change. The up, down, and unchanged loops identified in this analysis were used for downstream analysis. To remove any potential biases, we resized the loop anchors to 10 kb by calculating the mid coordinates and expanding 5 kb downstream and 5 kb upstream. We used these loops coordinates to perform aggregated peak analysis on the Hi-C contact matrices using GENOVA (van der Weide et al., 2021) with default options. All enrichment analysis of PCMC loops with different data sets with genomic features were performed using bedtools (Quinlan & Hall, 2010).

## Results

### A/B compartments in mES cells are resilient to BRG1 or INO80 depletion

To investigate the role of the esBAF and INO80C nucleosome remodeling complexes in shaping the three-dimensional (3D) genome architecture of mouse embryonic stem (mES) cells, we performed 48 h esiRNA-mediated knockdown of *Smarca4* (encoding BRG1) or *Ino80*, the catalytic subunits of esBAF and INO80C, respectively (**Fig. S1A**). *In situ* Hi-C was subsequently performed to assess genome-wide chromatin interactions. For each condition, two biological replicates were combined, yielding 1,181,605,537, 1,205,081,983, and 1,180,977,780 valid Hi-C contacts for control, *Smarca4* knockdown (KD), and *Ino80* KD samples, respectively (**Figs. 1A-C** and **S1B-D**). These combined datasets achieved a resolution of 2 kb as defined previously (Rao et al., 2014).

**Fig 1.**
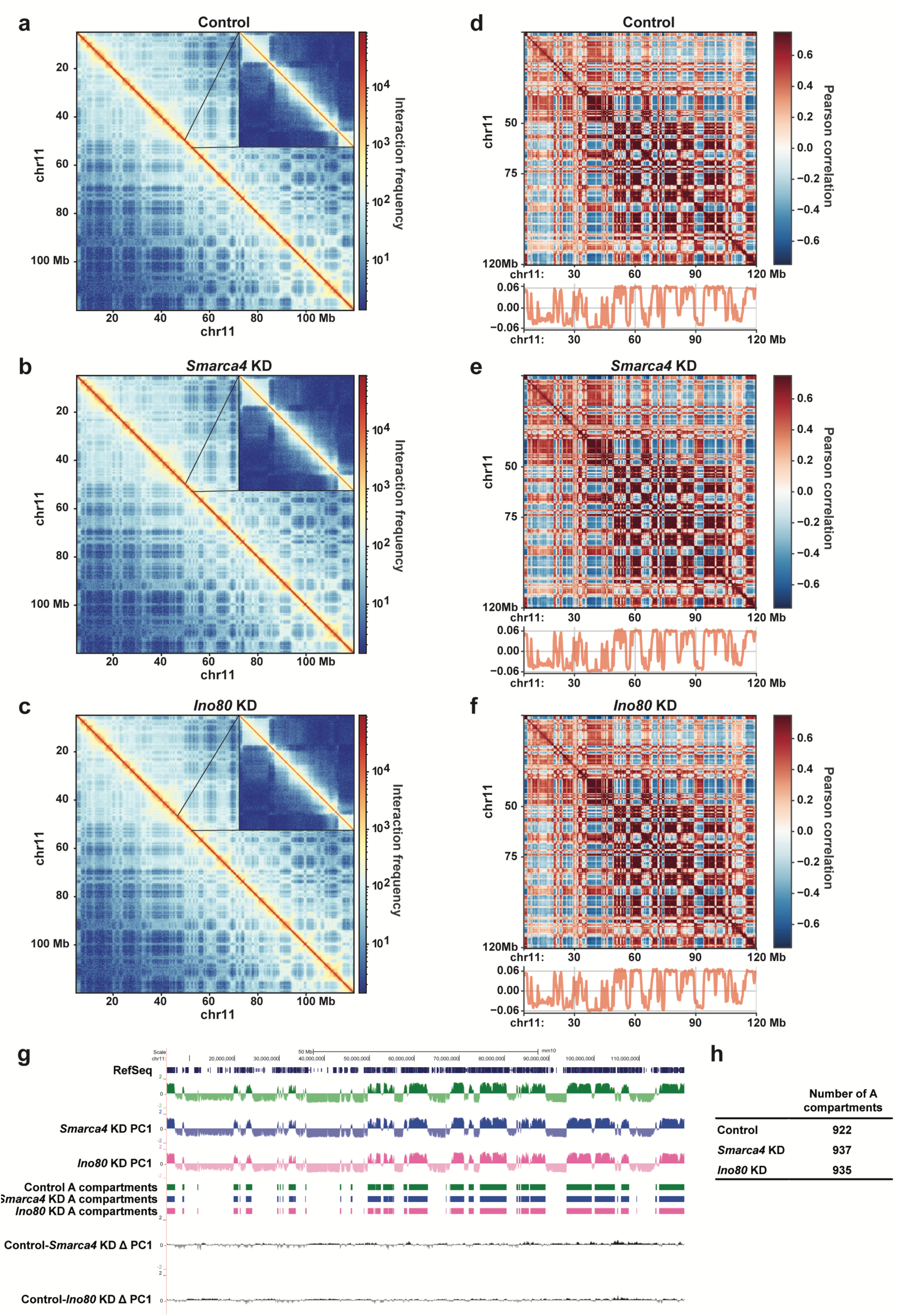
Depletion of *Smarca4* or *Ino80* does not alter A/B compartments in mES cells. **a-c)** Knight-Ruiz (KR) (Knight & Ruiz, 2013) balanced log1p interaction map for chromosome 11 at 250 kb resolution for the combined replicates for control sample (a), *Smarca4* KD (b), and *Ino80* KD (c). Right corner is a zoom-in. The interaction maps were created using HiCExplorer (Wolff et al., 2020). **d-f)** Pearson correlation maps at 250 kb for chromosome 11 of the combined interaction for control (d), *Smarca4* KD (e), and *Ino80* KD (f). The correlation maps were generated using FAN-C (Kruse et al., 2020). **g)** UCSC genome browser showing PC1 values and A compartments (obtained with HOMER), corresponding to control (green), *Smarca4* KD (blue), and *Ino80* KD (pink). *Smarca4* KD over control and *Ino80* KD over control differential PC1 values are shown below. **h)** Number of A compartments for each combined dataset.

Given that A/B compartments broadly correspond to euchromatic and heterochromatic regions and can change during differentiation and between cell types (Dixon et al., 2015; Lieberman-Aiden et al., 2009), we considered whether loss of nucleosome remodelers might perturb compartmental organization. Because remodelers are generally thought to act locally at genes and enhancers, any compartment-scale effects would most plausibly be indirect—for example, mediated by altered cell state or cell-cycle distribution following remodeler loss. To test this, we generated genome-wide Pearson correlation matrices with FAN-C (Kruse et al., 2020) (**Figs. 1D,F** and **S1E-G**), and performed principal component analysis at 25 kb resolution using HOMER (Heinz et al., 2010) to define A/B compartments (**Figs. 1G** and **S1HG**).

Comparison of PC1 signals between knockdown and control conditions revealed only minimal changes: the number and genomic positions of A compartments were largely conserved (**Figs. 1G,H** and **S1H**), and compartment profiles were reproducible across biological replicates (**Fig. S2**). Thus, in asynchronous mES cell populations, the proliferation and cell-cycle perturbations caused by BRG1 or INO80 depletion were not sufficient to induce detectable shifts compartment strength. We therefore conclude that short-term loss of these remodelers has predominantly local effects on chromatin architecture, though we cannot exclude compartment changes under synchronized or differentiation conditions.

### esBAF and INO80C maintain active subcompartments

Given the established roles of BRG1 and INO80 in transcription regulation and nucleosome remodeling (Ahmad et al., 2024; Bao & Shen, 2007; Klein & Hainer, 2020; Patty & Hainer, 2020; Poli et al., 2017; Willhoft & Wigley, 2020), we investigated whether depletion of *Smarca4* or *Ino80* alters genome subcompartment organization in mES cells. We applied CALDER2-based analysis to Hi-C datasets (Liu et al., 2021), which enables classification of genomic regions beyond the two-state model established by A/B compartmentalization (Kalluchi et al., 2023; Rao et al., 2014), capturing a spectrum of chromatin states defined by transcriptional activity, histone posttranslational modifications, and replication timing (MacKay & Kusalik, 2020). CALDER2 segments the genome into eight subcompartments, ranked from most open/active to most closed/inactive: A.1.1, A.1.2, A.2.1, A.2.2, B.1.1, B.1.2, B.2.1, and B.2.2.

We observed a modest increase in the total amount of the genome assigned to inactive subcompartments (B.1.1-B.2.2) upon either *Smarca4* or *Ino80* KD, consistently across biological replicates (**Fig. 2A**). Furthermore, more regions transitioned to a closed subcompartment state than to a more open state following depletion of either remodeler (**Figs. 2B** and **S3A**). Specifically, 15% (797/5155) and 11% (598/5221) of regions became more open, while 21% (1032/5155) and 17% (862/5221) became more closed in *Smarca4* and *Ino80* KD, respectively. These transitions were generally incremental, often involving neighboring subcompartments (e.g. A.2.1 to A.2.2 or A.1.2; **Fig. S3B,C**), but together suggest that esBAF and INO80C are required to preserve chromatin in an active state.

**Figure 2.**
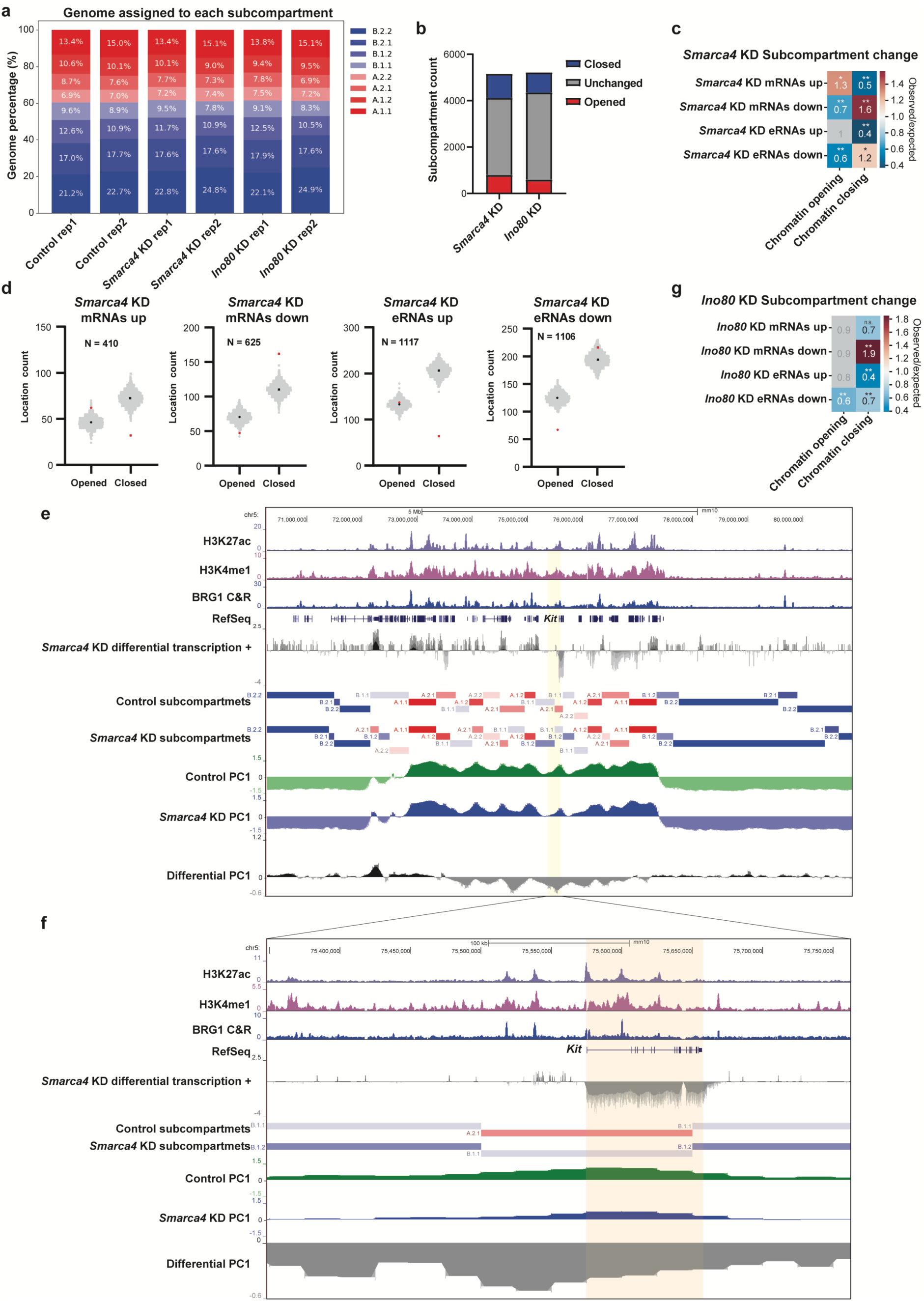
Depletion of *Smarca4* or *Ino80* leads to subcompartment changes. **a)** Proportion of genome assigned to open subcompartments (red: A.1.1, A.1.2, A.2.1, A.2.2) or closed subcompartments (blue: B.1.1, B.1.2, B.2.1, B.2.2) in control, *Smarca4* KD, or *Ino80* KD Hi-C replicate datasets. **b)** Number of subcompartment changes in *Smarca4* KD or *Ino80* KD Hi-C datasets relative to control. **c)** Observed over expected ratio of subcompartment changes in *Smarca4* KD relative to control Hi-C datasets at specific genomic locations (see Methods). Genomic features include all mRNAs with increased transcription or decreased transcription upon *Smarca4* KD relative to control TT-seq data (*Smarca4* mRNAs up and *Smarca4* KD mRNAs down (Patty et al., 2025)) and all putative eRNAs with increased transcription or decreased transcription from *Smarca4* KD relative to control TT-seq data (*Smarca4* eRNAs up and *Smarca4* KD eRNAs down). **d**) Count of subcompartment switches (openings or closing) for the corresponding locations; experimental value (red), 1,000 randomized location iterations (grey), and average of randomized locations (black). **e)** UCSC genome browser track for 10 Mb on chromosome 5 depicting H3K27ac and H3K4me1 ChIP-seq data (Chronis et al., 2017), BRG1 CUT&RUN, *Smarca4* KD relative to control TT-seq data, control and *Smarca4* KD subcompartment Hi-C data, and control, *Smarca4* KD and differential compartment Hi-C data. **f)** Zoom in of *Kit* locus on chromosome 5 with data as in e. **g)** As in C but for *Ino80* KD relative to control Hi-C datasets and using *Ino80* KD TT-seq data (Patty et al., 2025).

To determine whether these subcompartment transitions are functionally linked to gene regulation, we analyzed nascent transcriptomic (TT-seq) data (Patty et al., 2025). Genes upregulated upon *Smarca4* KD exhibited significantly increased chromatin opening and reduced chromatin closing than expected by chance, while downregulated genes showed the opposite trend (**Figs. 2C** and **S3D**). Similar patterns were observed for putative enhancer RNAs (eRNAs; **Figs. 2C** and **S3D**). These shifts were validated by comparing the number of observed versus expected transitions between open and closed states (**Fig. 2D**).

To assess whether subcompartment change is associated with BRG1 occupancy, we performed CUT&RUN profiling of BRG1 (**Fig. S3E**). Genomic regions bound by BRG1 exhibited significantly reduced chromatin opening following *Smarca4* KD compared to control. These effects were consistent, albeit less pronounced, when using previously published ChIP-seq data (Ho et al., 2009); **Fig. S3F-H**). At transcription start sites (TSSs), BRG1-bound promoters showed a marked decrease in compartment opening relative to unbound promoters (compare TSS BRG1 to TSS no BRG1; **Fig. S3F-H**). A similar decrease in chromatin opening was observed at distal DNase I hypersensitive sites (DHSs) occupied by BRG1, suggesting that BRG1 maintains local chromatin architecture at regulatory elements (compare DHS BRG1 to DHS no BRG1; **Fig. S3F-H**). These results further indicate that BRG1 binding has a role in maintaining the proper state of chromatin locally.

As an example, the *Kit* locus exhibited a subcompartment shift from A.2.1 to B.1.1 following *Smarca4* KD (**Figs. 2E,F** and **S3I**). Subcompartment shifts were reproducible (**Figs. S3A,D,G,I; S4A,D and S5**). Consistent with this shift, *Kit* was downregulated in the mES cell TT-seq data (**Fig. 2E,F**). Visualization of publicly available H3K4me3 and H3K27ac ChIP-seq datasets (Chronis et al., 2017) and BRG1 CUT&RUN datasets show BRG1, H3K4me3, and H3K27ac enrichment at the *Kit* promoter and upstream enhancer regions, supporting direct regulatory involvement. Subcompartment closure at *Kit* was corroborated by decreased PC1 values in the *Smarca4* KD using traditional A/B compartment assignment using principal component analysis of Hi-C data across biological replicates (**Fig. 2E,F**).

Compared to *Smarca4* depletion, the effects observed upon *Ino80* depletion in subcompartment rearrangements were more limited to the loci where transcription is affected by *Ino80* loss. Specifically, we observed strong subcompartment closing at genes downregulated in *Ino80* KD TT-seq data (**Figs. 2G** and **S4A,B**). Although chromatin opening was not enriched at either promoters or enhancers with increased transcription in *Ino80* KD, we observed reduced subcompartment closing relative to expected (**Figs. 2G** and **S4A,B**). To further probe this relationship, we utilized high-quality INO80 ChIP-seq data (Xue et al., 2017). While no global trend was detected at all INO80-bound sites, TSSs and DHSs occupied by INO80 showed a significant decreased for subcompartment closing (**Fig. S4C-E**). One example is the *Zfp827* locus, which encodes a zinc finger transcription factor involved in gene regulation (Conomos et al., 2014; Hollensen et al., 2020; Sahu et al., 2022). In *Ino80* KD cells, *Zfp827* transcription was reduced and its subcompartment shifted from A.1.2 to A.2.1, consistent with a more closed chromatin state, also observed by traditional PC analysis (**Fig. S5**). Collectively, these results demonstrate that BRG1 and INO80 contribute to the maintenance of subcompartment organization, with effects that are especially pronounced at loci where these remodelers bind and/or their loss impacts transcription.

### esBAF and INO80C depletion do not alter TAD organization in mES cells

We next assessed whether depletion of *Smarca4* or *Ino80* affects TAD organization in mES cells. TADs were identified using HiCExplorer (Ramírez et al., 2018; Wolff et al., 2018, 2020), yielding 6,360, 6,421, and 6,174 domains in control, *Smarca4* KD, and *Ino80* KD datasets, respectively. The majority of TADs were conserved across conditions, with only 200 (∼3%) and 320 (∼5%) differential TADs in the *Smarca4* and *Ino80* KD datasets, respectively, when compared to control (**Figs. 3A** and **S6A**). To validate these findings using an independent method, we employed TADcompare (Cresswell & Dozmorov, 2020), which detects TAD boundaries in two datasets and categorizes them as either differential or non-differential. Consistent with the HiCExplorer-based analysis, the vast majority of TAD boundaries were classified as non-differential by TADcompare (**Figs. 3B** and **S6B,C**).

**Figure 3.**
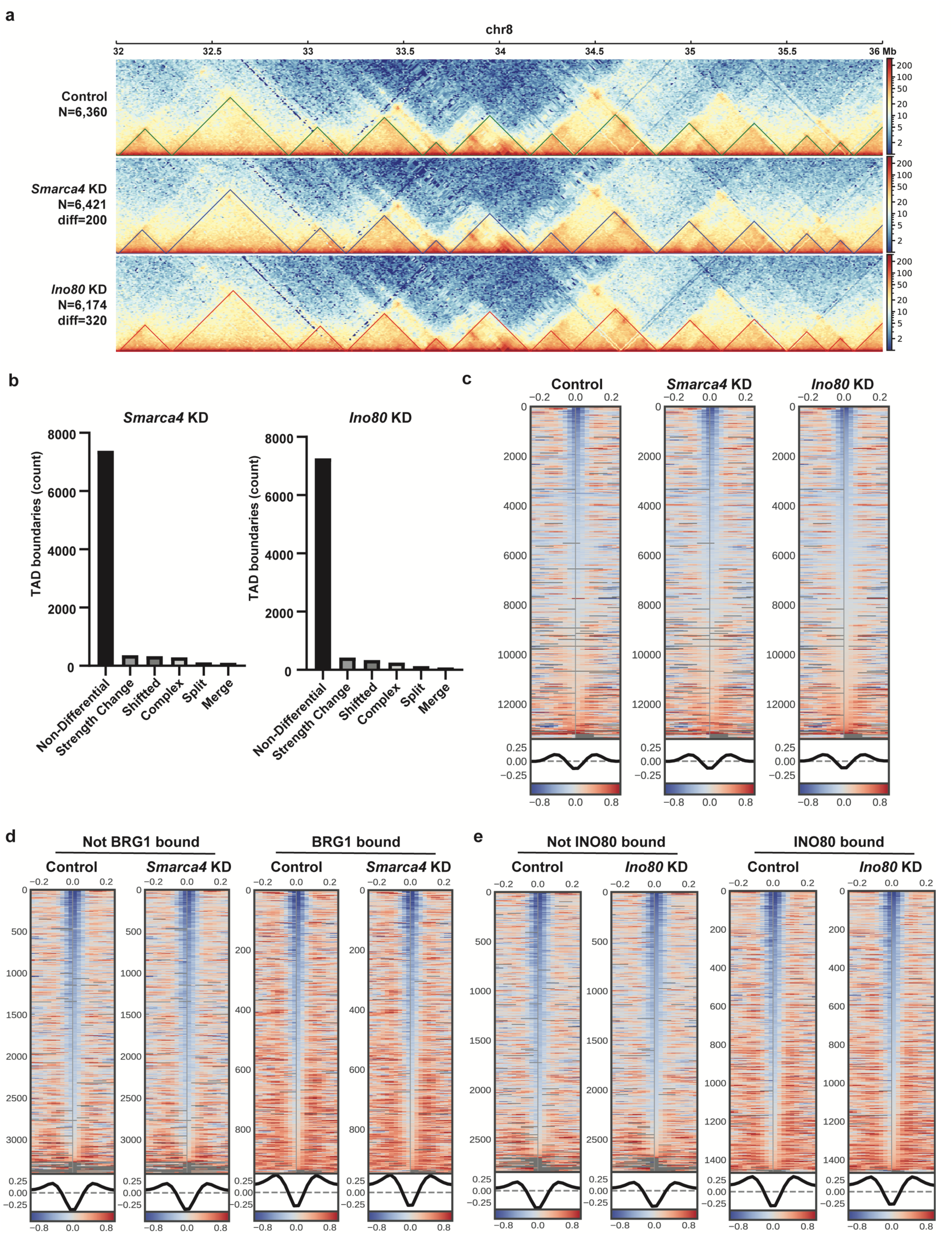
Depletion of *Smarca4* or *Ino80* does not alter TADs in mES cells. **a)** TAD interaction frequencies generated using HiCExplorer at 10 kb resolution over a 4 Mb portion of chromosome 8 for control (top) *Smarca4* KD (middle) and *Ino80* KD (bottom) merged replicas. **b)** Number of differential TAD boundaries in *Smarca4* KD (left) or *Ino80* KD (right) relative to control. **c)** Heatmap depicting insulation score over all TAD boundaries for control (left) *Smarca4* KD (middle) and *Ino80* KD (right) merged replicates generated using cooltools. **d)** Heatmap depicting insulation score over BRG1 unbound (left) or bound (right) TAD boundaries for control and *Smarca4* KD merged replicas generated using cooltools. **e)** Heatmap depicting insulation score over INO80 unbound (left) or bound (right) TAD boundaries for control and *Ino80* KD merged replicas generated using cooltools.

We further evaluated TAD structure by calculating insulation scores, a complementary approach that quantifies boundary strength. Using cooltools (Open2C et al., 2024), we found no significant changes in aggregate insulation scores in either *Smarca4* KD or *Ino80* KD datasets relative to control (**Figs. 3C** and **S6D,E**). Notably, this result held true even when restricting the analysis to TAD boundaries bound by BRG1 or INO80, indicating that loss of these remodelers does not affect boundary insulation strength at their binding sites (**Fig. 3D,E**). Together, these findings suggest that depletion of *Smarca4* or *Ino80* does not disrupt global TAD organization in mES cells. This is consistent with previous observations that BRG1 loss does not alter TAD structure in mES cells (Barisic et al., 2019), though it contrasts with a report from another cell type, MCF-10A, where BRG1 depletion was found to impact TAD boundaries (Barutcu et al., 2016).

### Hi-C loops are enriched for architectural features

To evaluate the potential role of esBAF and INO80C in regulating enhancer-promoter looping, we performed loop calling on merged Hi-C datasets at 10 kb resolution. We identified 5,903, 6,013, and 5,485 high-confidence chromatin loops in control, *Smarca4* KD, and *Ino80* KD conditions, respectively, with representative examples of conserved and differential loops shown in **Fig. 4A**. Specifically, we identified 1099 differential loops in *Smarca4* KD and 835 differential loops in *Ino80* KD. Notably, neither conserved nor differential loops exhibited significant enrichment for BRG1 or INO80 binding (**Fig. 4B,C**). However, both unchanged and differential loops in *Smarca4* KD and *Ino80* KD datasets were enriched for CTCF, consistent with known architectural features, with no enrichment for TSS-distal DHSs or H3K27ac in differential loops (**Fig. S7A,B**). These observations suggest that the subset of differential loops confidently identified in our Hi-C datasets primarily represent structural or architectural loops, rather than enhancer-promoter interactions.

**Figure 4.**
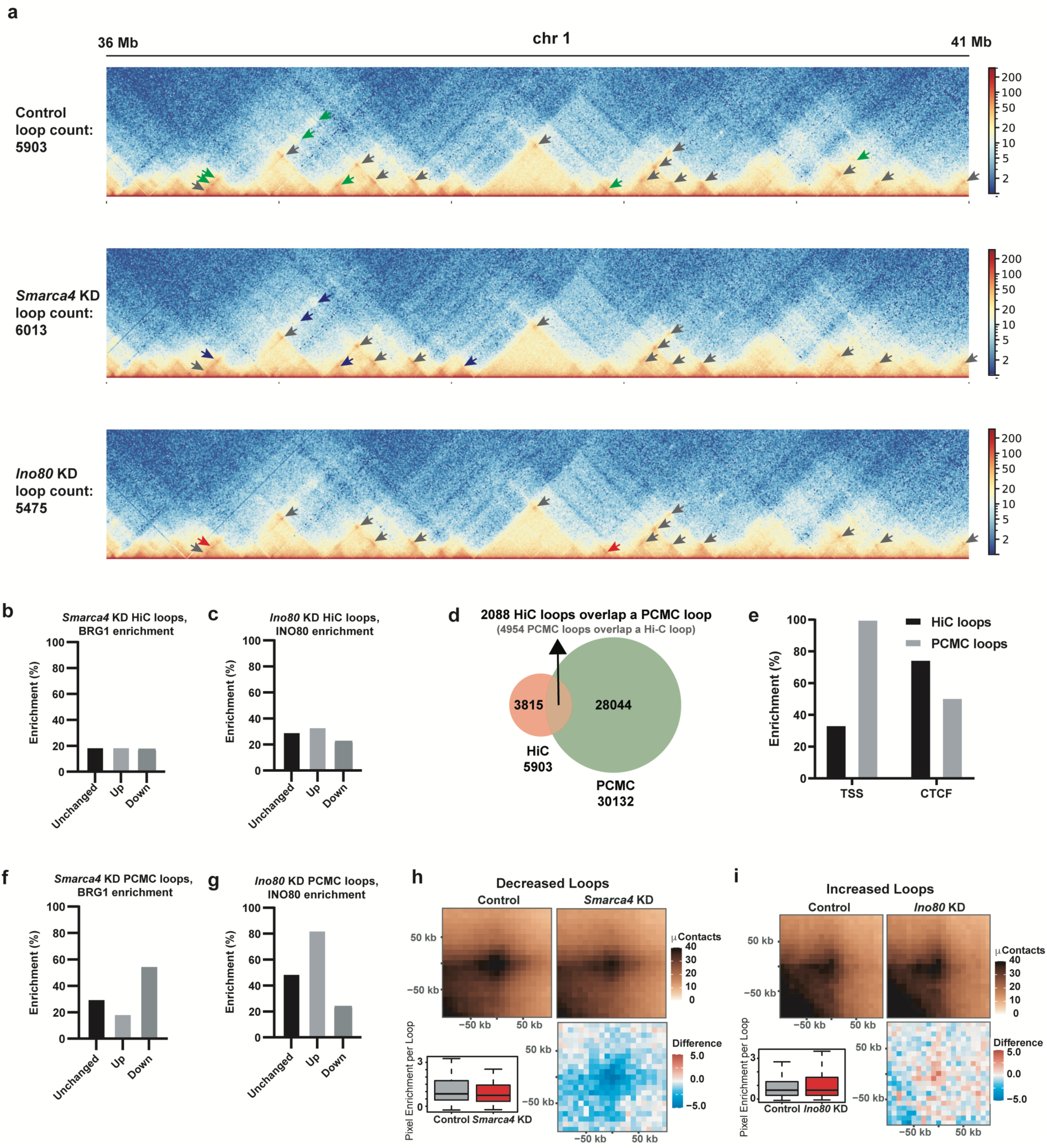
PCMC identifies differential enhancer-promoter loops in *Smarca4* and *Ino80* KD. **a)** Interaction frequencies generated using HiCExplorer at 10 kb resolution over a 5 Mb portion of chromosome 1 for control (top) *Smarca4* KD (middle) and *Ino80* KD (bottom) merged replicas. **b)** Percentage of unchanged, increased, or decreased loops in *Smarca4* KD Hi-C datasets relative to control that are also locations bound by BRG1 (from CUT&RUN) **c)** Percentage of unchanged, increased, or decreased loops in *Ino80* KD Hi-C datasets relative to control that are also locations bound by of INO80 (from ChIP-seq) (Xue et al., 2017) **d)** Overlap of loops called from control Hi-C or control PCMC datasets. The total number of loops in each dataset is written below the venn diagram. **e)** Enrichment of either loop anchor in Hi-C (black) or PCMC (PCMC; grey) control datasets over TSS locations or CTCF binding sites. **f)** Percentage of unchanged, increased, or decreased loops in *Smarca4* KD PCMC loops relative to control that are also locations bound by BRG1 (from CUT&RUN). **g)** Percentage of unchanged, increased, or decreased loops in *Ino80* KD PCMC loops relative to control that are also locations bound by INO80 (from ChIP-seq) (Xue et al., 2017). **h)** Aggregate peak analysis (APA) for decreased PCMC loops in *Smarca4* KD (top right) relative to control (top left). Quantified peak enrichment (bottom left) and differential interaction frequency signal (bottom right) are also shown. **i)** As in h for increased loops in *Ino80* KD.

### Promoter capture micro-C identifies differential enhancer-promoter loops in *Smarca4* and *Ino80* KD

Given the limited detection of enhancer-promoter loops in standard Hi-C datasets, we employed promoter capture micro-C (PCMC), utilizing three probes per promoter to enrich for promoter-anchored interactions. Efficient depletion of *Smarca4* or *Ino80* (∼90%) was achieved (**Fig. S7C,D**) and PCMC was performed in two biological replicates per condition. We identified ∼32,000 loops in each dataset with 399 differential loops in *Smarca4* KD and 450 differential loops in *Ino80* KD relative to control. As noted above, Hi-C-identified differential loops were predominantly CTCF-enriched (∼75%) and included only ∼25% TSS-based loops (**Fig. 4E**), consistent with prior studies (4D Nucleome Consortium et al., 2024). In contrast, 100% of loops detected via PCMC were TSS-anchored, as expected, with only ∼50% showing CTCF enrichment (**Figs. 4E** and **S7E,F**). These results are consistent with PCMC providing more direct access to functional enhancer-promoter contacts.

Functionally, reduced loops in *Smarca4* KD datasets were enriched for BRG1 binding (**Fig. 4F**) and overlapped with TSS-distal DHSs and H3K27ac peaks—hallmarks of active enhancers (**Fig. S7E**). In contrast, unchanged or increased loops in *Smarca4* KD did not show this enrichment. Loops that increased upon *Ino80* depletion were enriched for INO80 binding (**Fig. 4G**), TSS-distal DHSs, and H3K27ac (**Fig. S7F**), whereas decreased and unchanged loops were not enriched for these features. Aggregate Peak Analysis (APA) revealed visible reduction in loop strength for *Smarca4* KD-sensitive loops and a visible gain in loop strength upon *Ino80* KD (**Fig. 4H,I**). In contrast, loops categorized as unchanged or those increased (*Smarca4* KD) or decreased (*Ino80* KD) showed little change in APA signal (**Fig. S7G,H**). Collectively, these findings indicate that esBAF promotes a subset of enhancer-promoter loops, while INO80C restricts or destabilizes specific loops. These effects appear to be context-dependent and most evident at regions bound by the respective nucleosome remodeler complexes.

### Differential PCMC loops are enriched for bivalent locations

To better understand the nature of the PCMC altered loops, we examined binding of pluripotency-related transcription factors over unchanged, increased, or decreased loops using publicly available CUT&RUN datasets (Hainer et al., 2019) (**Fig. 5A,B**). Both BRG1 and INO80 binding in mES cells overlaps with pluripotency-related factors, including OCT4, SOX2, and NANOG (OSN) (Hainer & Fazzio, 2015; King & Klose, 2017; L. Wang et al., 2014). Loops decreased in *Smarca4* KD or increased in *Ino80* KD were enriched for OSN, reflecting loops that are also enriched for BRG1 and INO80, respectively (**Fig. 4F,G**).

**Figure 5.**
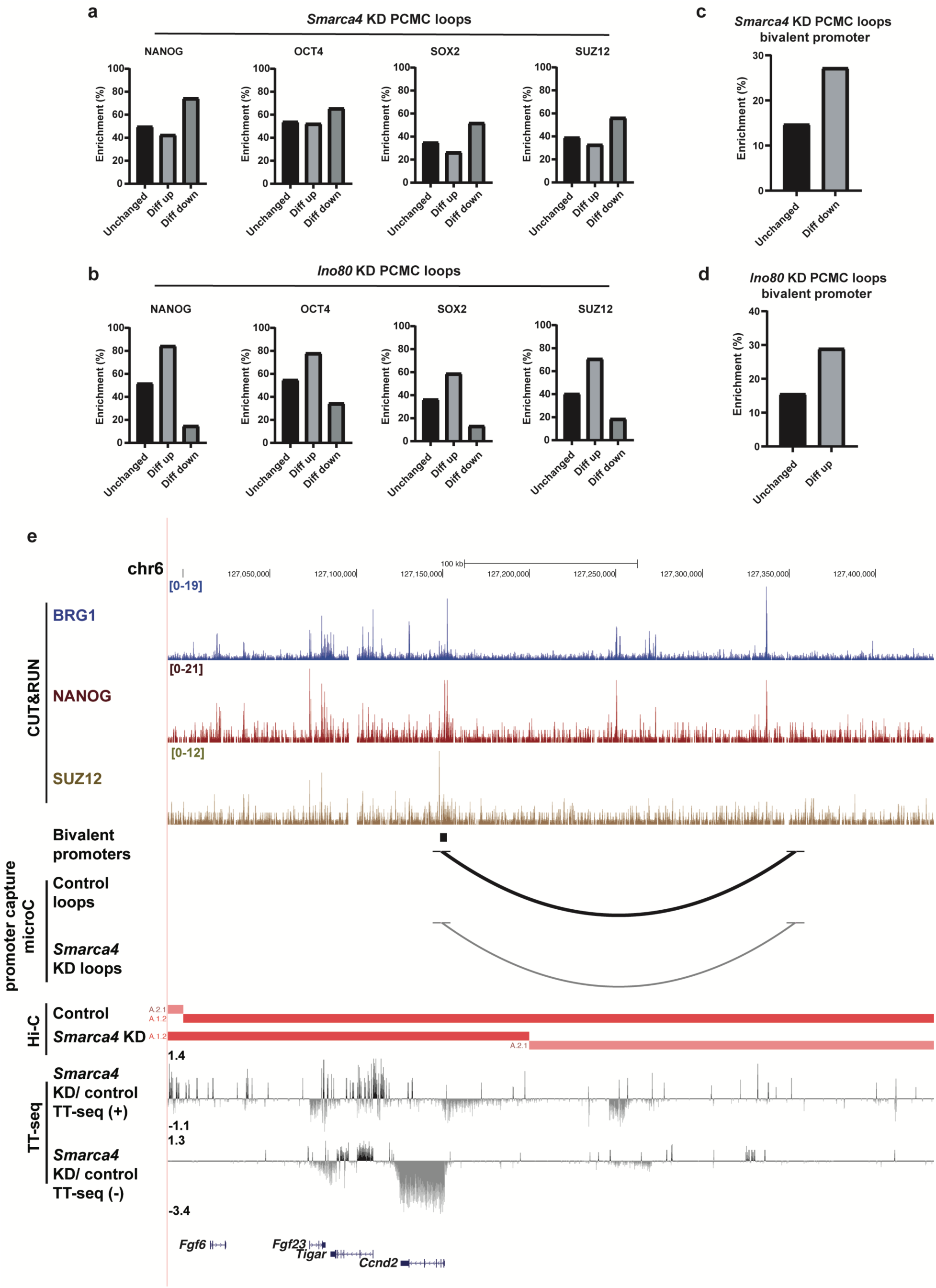
Differential PCMC loops are enriched for bivalent locations bound by pluripotency factors. **a)** Percentage of unchanged, increased, or decreased loops in *Smarca4* KD PCMC loops relative to control also bound by OSN or SUZ12 (Hainer et al., 2019). **b)** As in a, for *Ino80* KD PCMC loops. **c)** Percentage of unchanged or decreased loops in *Smarca4* KD PCMC loops characterized as bivalent (Seneviratne et al., 2024). **d)** Percentage of unchanged or increased loops in *Ino80* KD PCMC loops characterized as bivalent (Seneviratne et al., 2024). **e)** UCSC genome browser track for 0.5 Mb on chromosome 6 depicting BRG1, NANOG, and SUZ12 CUT&RUN, bivalent promoter calls (Seneviratne et al., 2024), PCMC loops in control and *Smarca4* KD, and compartment calls from Hi-C data in control and *Smarca4* KD.

Previous studies found that these pluripotency factors bind a subset of bivalent genomic locations enriched for PRC2 (Boyer et al., 2006; Heurtier et al., 2019; Shan et al., 2017; Squazzo et al., 2006). Furthermore, BRG1 acts in a context specific manner with PRC2 where the BAF complex can either cooperate with PRC2 to promote repression (Alexander et al., 2015; Ho et al., 2011; Semrau et al., 2017; Weber et al., 2021) or antagonize PRC2 to promote activation (Braun et al., 2017; Kadoch et al., 2017; Kennison & Tamkun, 1988; Shao et al., 1999; B. G. Wilson et al., 2010; M. R. Wilson et al., 2022) whereas INO80C helps to recruit PRC2 (Chakraborty & Magnuson, 2022; Yu et al., 2021). Therefore, we examined whether loops altered in *Smarca4* or *Ino80* KD were enriched for PRC2 by analyzing publicly available SUZ12 localization data (Hainer et al., 2019) (**Fig. 5A,B**). Similar to OSN, loops decreased in *Smarca4* KD or increased in *Ino80* KD were enriched for SUZ12. To further examine the characteristics of these locations, we intersected the loop coordinates with previously called bivalent locations (Seneviratne et al., 2024)(**Fig. 5C,D**). Loops decreased in *Smarca4* KD or increased in *Ino80* KD were more frequently enriched for bivalent locations, suggesting that the loops sensitive to loss of these remodelers are bivalent locations enriched for pluripotency factors. One example is shown for *Ccnd1* which is bound by BRG1, NANOG, and SUZ12 and identified as a bivalent promoter (**Fig. 5E**). The loop identified in control PCMC datasets is reduced upon *Smarca4* depletion, as is transcription of *Ccnd1* and a subcompartment switch is observed (**Fig. 5E**). Together, these data suggest that the enhancer-promoter loops esBAF promotes and INO80C restricts are pluripotency and bivalency related.

## Discussion

Our study reveals that the nucleosome remodelers esBAF and INO80, while extensively characterized for their roles in local chromatin accessibility and transcriptional regulation, have limited influence on large-scale 3D genome architecture. Specifically, we find that global features such as A/B compartments and topologically associating domains (TADs) remain largely intact upon loss of either remodeler, suggesting that these higher-order domains are maintained independently of esBAF and INO80C activity (**Figs. 1** and **3**). However, at a finer resolution, we observe that subcompartment organization is sensitive to remodeler depletion, indicating a more nuanced and direct role for these complexes in shaping chromatin topology at sub-megabase scales (**Fig. 2**). This selective disruption of subcompartments may reflect localized changes in chromatin state or nuclear positioning that lead to changes in nearby gene expression but do not impact broader domain structures. To further investigate potential changes in gene regulatory architecture, we employed PCMC, which provides high-resolution detection of promoter-anchored chromatin interactions (**Fig. 4F,G**). This approach enabled us to identify a subset of promoter-anchored loops whose interaction frequencies were altered following loss of esBAF or INO80C. Although the specific loops affected differed between the two remodelers, they were frequently associated with bivalent chromatin regions co-bound by the core pluripotency factors OSN, as well as BRG1 and INO80 themselves (**Fig. 5A,B**). These findings suggest that esBAF and INO80C are not general regulators of chromatin looping but may instead fine-tune regulatory interactions at distinct key developmental loci in ES cells. Taken together, our results highlight a previously underappreciated role for nucleosome remodelers in modulating specific aspects of higher-order genome organization, particularly within regulatory domains critical for maintaining pluripotent identity. The specific domains regulated by remodelers are likely cell-type specific, with complexes performing alternative functions in different cell types, as observed in (Barutcu et al., 2016). By uncovering novel functions for nucleosome remodelers in higher-order genome folding, this work advances our understanding of how chromatin structure integrates with chromatin states and transcriptional networks to sustain stem cell identity and guide cell fate decisions.

The role of BRG1 in the formation and maintenance of TAD boundaries has been variably reported across different cellular contexts. In human mammary epithelial MCF-10A cells, BRG1 knockdown was associated with a modest weakening of TAD boundary strength, suggesting a potential architectural role for this remodeler in organizing 3D genome structure (Barutcu et al., 2016). In contrast, studies in mES cells employing inducible deletion of BRG1 observed no significant changes in TAD organization (Barisic et al., 2019), raising questions about whether BRG1’s contribution to TAD boundaries is context-dependent. To reconcile these conflicting reports, we conducted a detailed analysis of TAD boundaries in our system using high-resolution Hi-C. Our results revealed that TAD boundaries were largely preserved upon BRG1 depletion, with no substantial loss in boundary strength or insulation. These findings are in line with Barisic et al., 2019 and suggest that BRG1 is not a core determinant of TAD boundary formation in this context. One possibility is that BRG1’s role in genome architecture is limited to finer-scale chromatin features, such as subcompartments and/or regulatory loops, rather than broader domain boundaries. Alternatively, TAD boundaries may be maintained by BRG1 in only specific cell types. Together, our results support the idea that BRG1 plays little to no role in the maintenance of TAD boundaries in mES cells and highlight the importance of considering cell type-specific chromatin environments when interpreting the functions of nucleosome remodelers.

Subcompartments represent a more recently characterized and nuanced layer of higher-order chromatin organization, emerging as a refinement of the classical A/B compartment model. Unlike the binary A/B classification, which broadly distinguishes active and inactive chromatin, subcompartments are defined by more specific combinations of chromatin features, including histone post-translational modifications, transcriptional activity, and nuclear positioning. This finer-grained organization captures functionally distinct chromatin environments that are often linked to specific gene regulatory programs and cellular states. Given the close relationship between subcompartment identity and underlying chromatin state, it is perhaps not surprising that we found esBAF and INO80C exert a measurable impact on subcompartment structure. These remodelers are known to regulate nucleosome positioning and accessibility, functions that are intimately tied to histone modification patterns and transcriptional output, both of which contribute to subcompartment identity. Our observation that subcompartments are altered upon remodeler loss, even in the absence of global changes to compartments or TADs, suggests that nucleosome remodelers play a more targeted role in fine-tuning chromatin environments at sub-megabase scales. This highlights the potential for nucleosome remodelers to influence genome organization in a more localized and context-dependent manner, potentially through interactions with histone modifiers, transcription factors, or nuclear architectural components that help define subcompartment domains.

While our Hi-C datasets were deeply sequenced, they lacked sufficient depth to enable robust detection of enhancer-promoter interactions. Instead, most loops identified from these data likely represent stable, architectural loops anchored by structural proteins such as CTCF and cohesin. To overcome this limitation and more directly assess regulatory interactions, we utilized PCMC, which enriches for contacts anchored at gene promoters. Although depletion of BRG1 or INO80 resulted in altered interaction frequencies at a subset of these loops, the vast majority remained unchanged. This relative stability may reflect functional redundancy among nucleosome remodelers, a limited requirement for BRG1 or INO80 in maintaining established loops, or the robustness of enhancer-promoter interactions once established. Alternatively, residual protein following knockdown or the extended time frame of depletion (48 hours) may obscure more transient roles. These findings underscore the complexity of interpreting chromatin looping phenotypes and suggest that while esBAF and INO80C can modulate specific regulatory interactions, they are not broadly required for maintaining the global landscape of promoter-based looping.

Although the specific enhancer-promoter loops affected by BRG1 and INO80 depletion were largely distinct, they converged on a common chromatin context: bivalent domains co-occupied by the core pluripotency transcription factors OSN, as well as BRG1 and INO80 themselves. These bivalent regions are characterized by the simultaneous presence of activating (H3K4me3) and repressive (H3K27me3) histone modifications and are thought to poise developmental genes for rapid activation or repression during cell fate transitions. The co-localization of BRG1 or INO80 with OSN at these loci is consistent with previous reports showing that both remodelers are physically and functionally associated with pluripotency transcriptional networks (Hainer & Fazzio, 2015; King & Klose, 2017; L. Wang et al., 2014).

BRG1, as the ATPase subunit of esBAF, has been shown to facilitate chromatin accessibility at OSN-bound enhancers, while INO80C has been implicated in both nucleosome positioning and the resolution of transcription-replication conflicts at active genes. The enrichment of altered loops at OSN-bound, bivalent regions suggests that rather than globally maintaining chromatin loops, esBAF and INO80C may act more selectively to modulate enhancer-promoter communication at genes crucial for lineage specification. These regions are also frequently co-occupied by PRC2, which deposits H3K27me3 and reinforces transcriptional repression. Thus, the interplay between nucleosome remodeling by BRG1 and INO80, OSN-mediated transcriptional priming, and PRC2-mediated repression may collectively establish and modulate a poised chromatin state that supports pluripotency while maintaining developmental potential. Our findings point to a model in which esBAF and INO80C serve distinctively as regulatory fine tuners of 3D chromatin architecture at these key developmental hubs, rather than as global organizers of chromatin topology. In future studies, it will be interesting to investigate the importance of these remodeler-based interactions during development.

## Data Availability

Hi-C and promoter capture Micro-C (PCMC) data have been deposited at the Gene Expression Omnibus and are publicly available as of the date of publication. All other data supporting the key findings are available in the article and supplementary materials.

## Acknowledgements

We are grateful to Johan Gibcus for guidance and members of the Hainer, Dekker, and Fazzio labs for helpful discussions.

## Supplementary Figures

**Figure S1.**
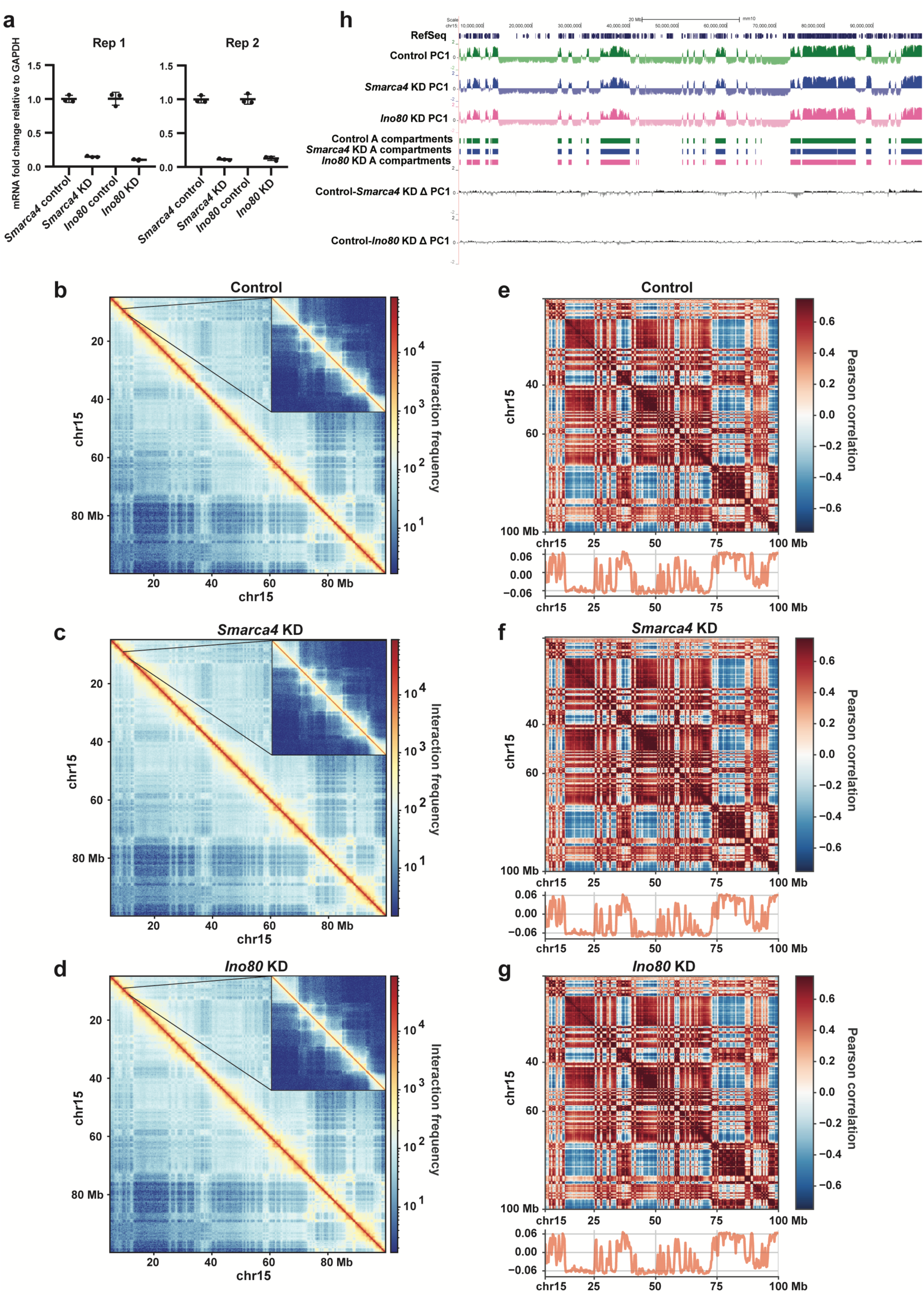
Depletion of *Smarca4* or *Ino80* does not alter A/B compartments at chromosome 15 in mES cells. **a)** RT-qPCR results showing mRNA fold change of *Smarca4* and *Ino80* in knockdown and control cells. Primers targeting *Smarca4* or *Ino80* were used (see Table S1). All values were normalized to *Gapdh* mRNA levels using the ΔΔCt method. Error bars represent ± standard deviation. Each biological replicate was measured in triplicate. **b-d)** Knight-Ruiz (KR) (Knight & Ruiz, 2013) balanced log1p interaction map for chromosome 15 at 250 kb resolution for the combined control sample (b), *Smarca4* KD (c), and *Ino80* KD (d). Right corner is a zoom-in. The interaction maps were created using HiCExplorer (Wolff et al., 2020). **e-g)** Pearson correlation map at 250 kb for chromosome 15 of the combined interaction for control (e), *Smarca4* KD (f), and *Ino80* KD (g). The correlation maps were generated using FAN-C (Kruse et al., 2020). **h)** UCSC genome browser showing PC1 values and A compartments corresponding to control (green), *Smarca4* KD (blue), and *Ino80* KD (pink). *Smarca4* KD over control and *Ino80* KD over control differential PC1 values are shown.

**Figure S2.**
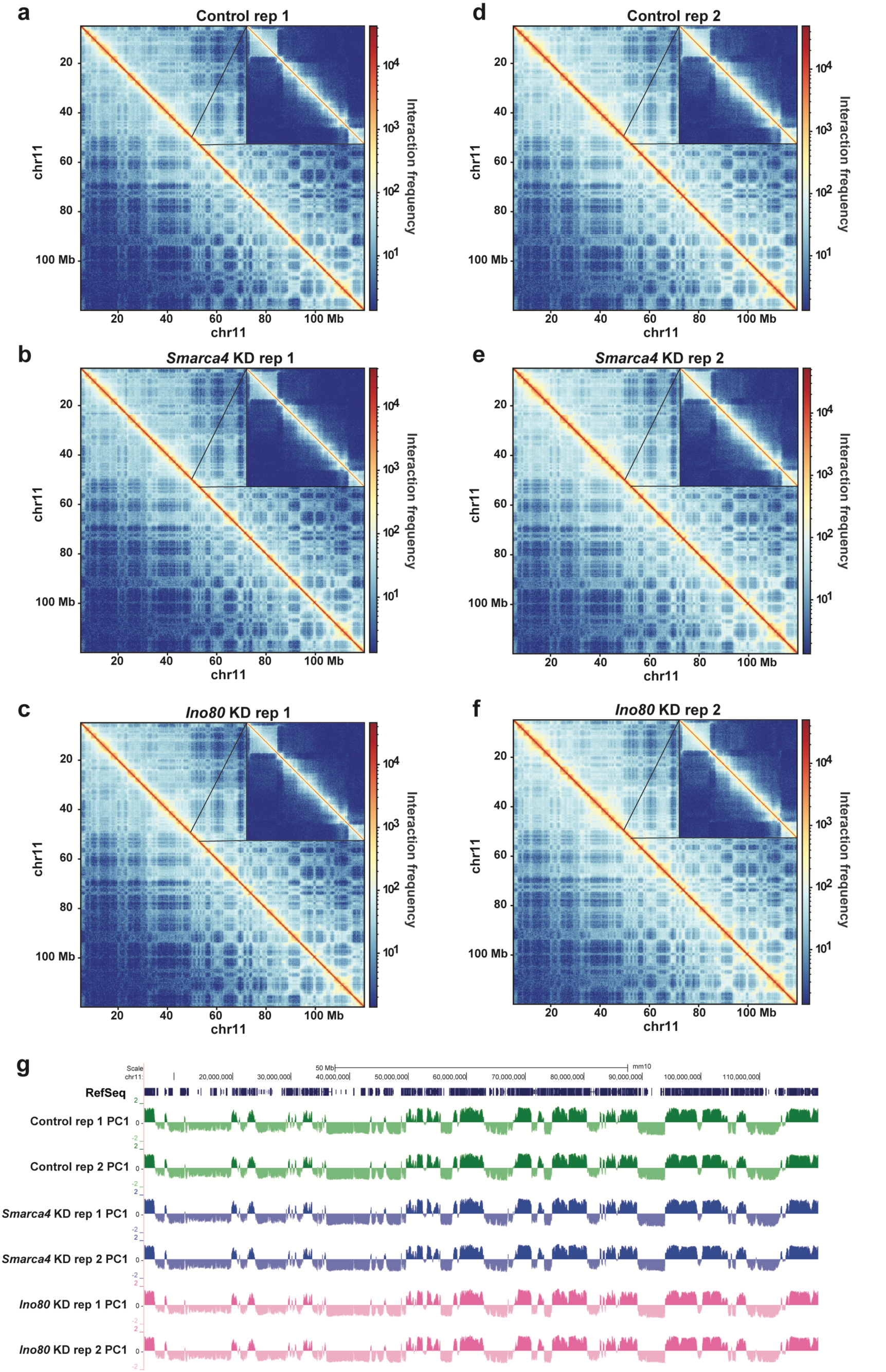
A/B compartments are reproducible across biological replicates. **a-c)** Knight-Ruiz (KR) (Knight & Ruiz, 2013) balanced log1p interaction map for chromosome 11 at 250 kb resolution for the one replicate of control (a), *Smarca4* KD (b), and *Ino80* KD (c), generated using HiCExplorer (Wolff et al., 2020). Right corner is a zoom-in. **d-f)** KR balanced log1p interaction map for chromosome 11 at 250 kb resolution for the second replicate of control (d), *Smarca4* KD (e), and *Ino80* KD (f), generated using HiCExplorer (Wolff et al., 2020). **g)** UCSC Genome browser showing A compartments identified using HOMER (Heinz et al., 2010), corresponding to the control (green), *Smarca4* KD (blue), and *Ino80* KD (pink) replicates.

**Figure S3.**
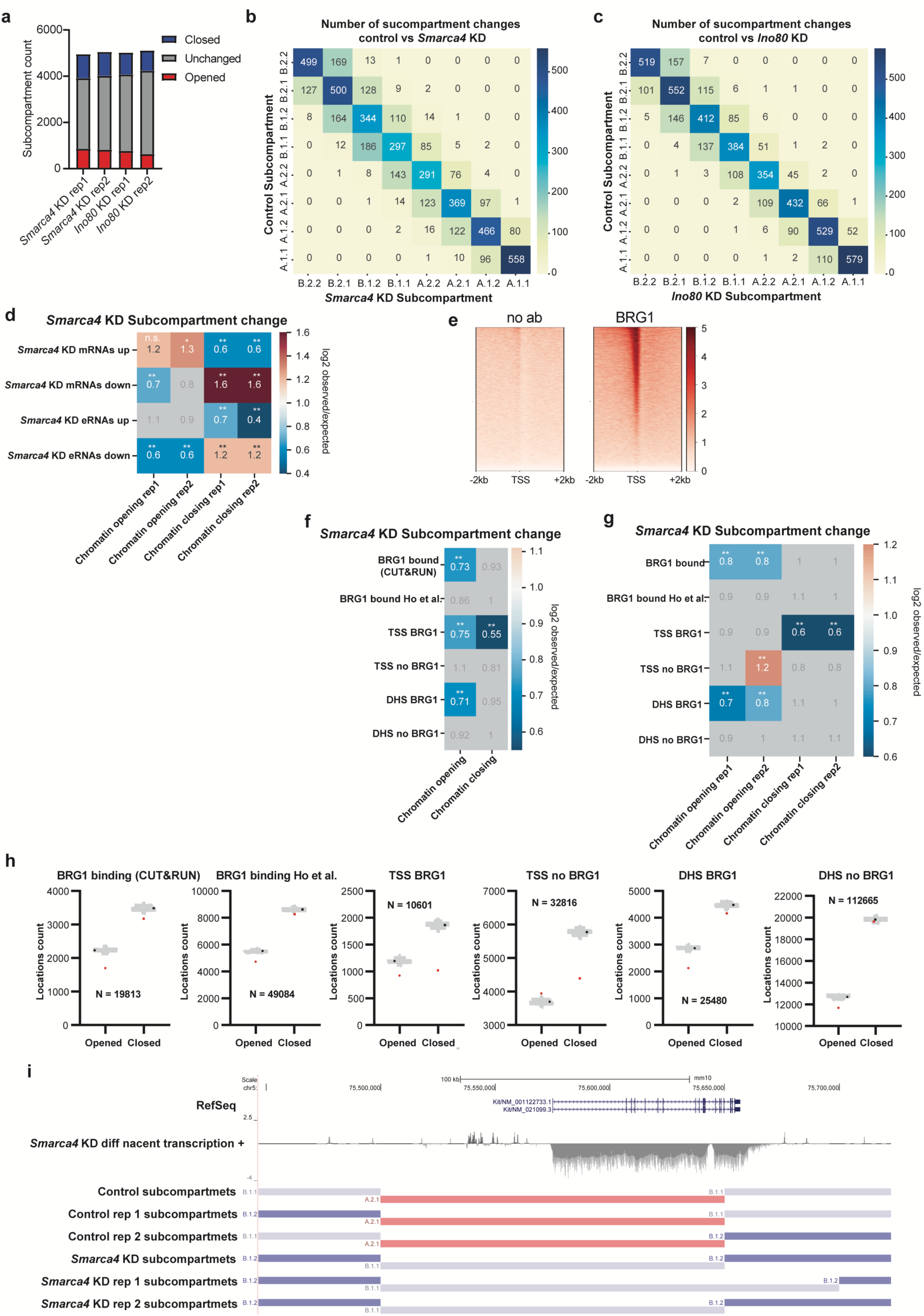
*Smarca4* KD leads to subcompartment changes relative to control Hi-C data. **a**) Number of subcompartment changes in *Smarca4* KD or *Ino80* KD Hi-C replicate datasets relative to control. **b-c**) Number of subcompartment changes for *Smarca4* KD (b) or *Ino80* KD (c) relative to control. **d**) Observed over expected ratio of subcompartment changes in *Smarca4* KD relative to control Hi-C dataset replicates at all mRNAs with increased transcription or decreased transcription from TT-seq data (Patty et al., 2025) (*Smarca4* mRNAs up and *Smarca4* KD mRNAs down) and all putative eRNAs with increased transcription or decreased transcription from *Smarca4* KD relative to control TT-seq data (*Smarca4* eRNAs up and *Smarca4* KD eRNAs down). **e**) Heatmaps of merged BRG1 CUT&RUN and negative control (noab) plotted over TSSs +/- 2 kb. **f**) Observed over expected ratio of subcompartment changes in *Smarca4* KD relative to control Hi-C dataset at all BRG1 bound locations from CUT&RUN data (BRG1 bound CUT&RUN); all BRG1 bound locations from previously published ChIP-seq data (BRG1 bound Ho et al); transcription start sites (TSSs) with BRG1 bound (from CUT&RUN data; TSS BRG1); TSSs with no BRG1 bound (TSS no BRG1); promoter distal DNase1 hypersensitive sites (DHSs) with BRG1 bound (DHS BRG1); promoter distal DHSs with no BRG1 bound (DHS no BRG1). **g)** As in f, but for biological replicates. **H)** Plots for subcompartment location counts in red for each genomic feature (as in b), with 1,000 randomized genomic locations independently generated shown in light grey and the average randomization shown in black. **i)** UCSC genome browser track for *Kit* locus as in Fig 2G, showing both merged and independent replicates for subcompartment calls.

**Figure S4.**
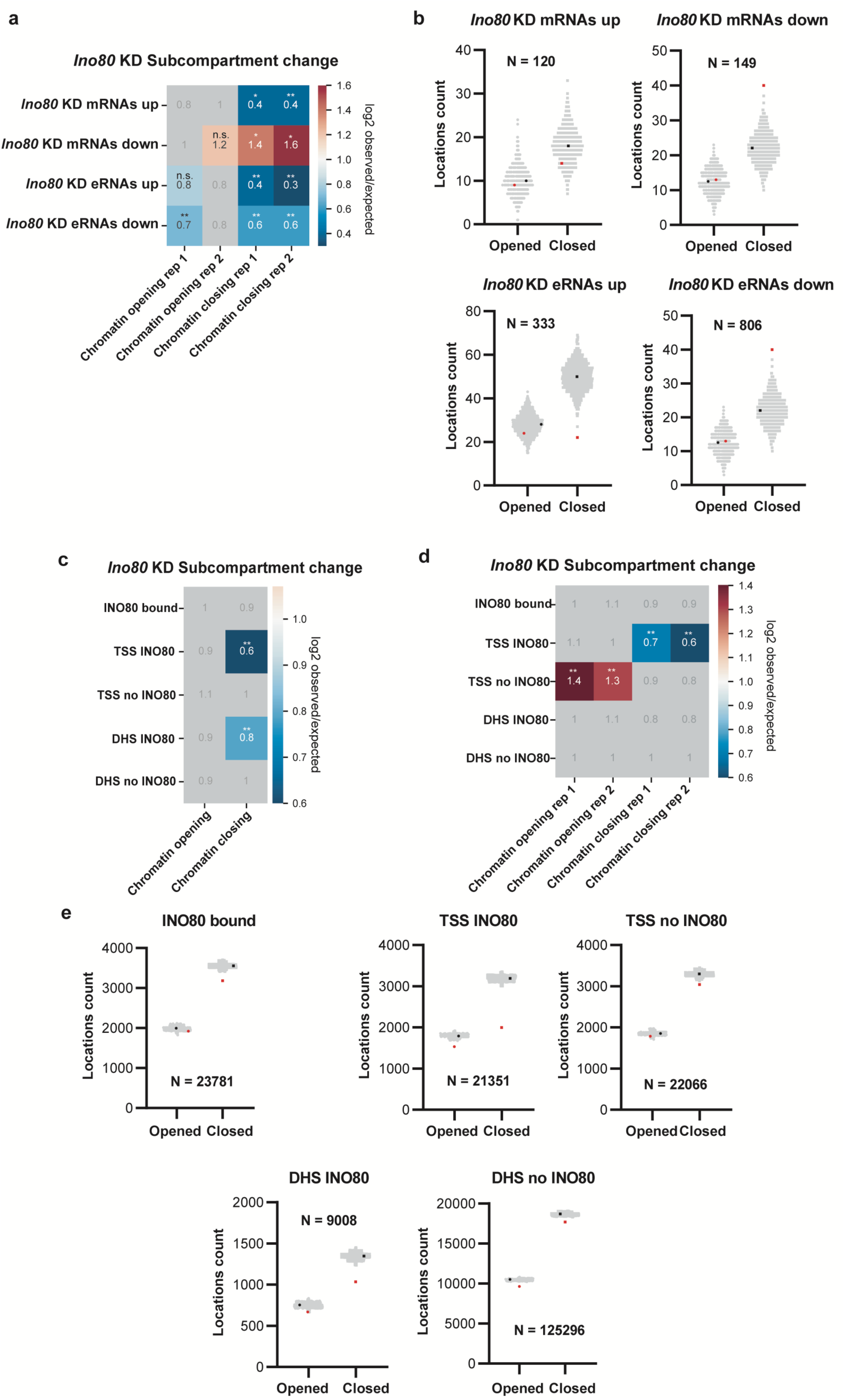
*Ino80* KD leads to subcompartment changes relative to control Hi-C data. **a)** Observed over expected ratio of subcompartment changes in *Ino80* KD relative to control Hi-C datasets for individual replicates at all mRNAs with increased transcription or decreased transcription from *Ino80* KD relative to control TT-seq data (*Ino80* mRNAs up and *Ino80* KD mRNAs down (Patty et al., 2025)) and all putative eRNAs with increased transcription or decreased transcription from *Ino80* KD relative to control TT-seq data (*Ino80* eRNAs up and *Ino80* KD eRNAs down). **b)** Plots for subcompartment location counts in red for each genomic feature (as in a), with 1,000 randomized genomic locations independently generated shown in light grey and the average randomization shown in black. **c)** Observed over expected ratio of subcompartment changes in *Ino80* KD relative to control Hi-C datasets at all INO80 bound locations from published ChIP-seq data (INO80 bound) (Xue et al., 2017); transcription start sites (TSSs) with INO80 bound (TSS INO80); TSSs with no INO80 bound (TSS no INO80); promoter distal DNase1 hypersensitive sites (DHSs) with INO80 bound (DHS INO80); promoter distal DHSs with no INO80 bound (DHS no INO80). **e)** As in C, but for individual replicates. **e)** Plots for subcompartment location counts in red for each genomic feature (as in c), with 1,000 randomized genomic locations independently generated shown in light grey and the average randomization shown in black.

**Figure S5.**
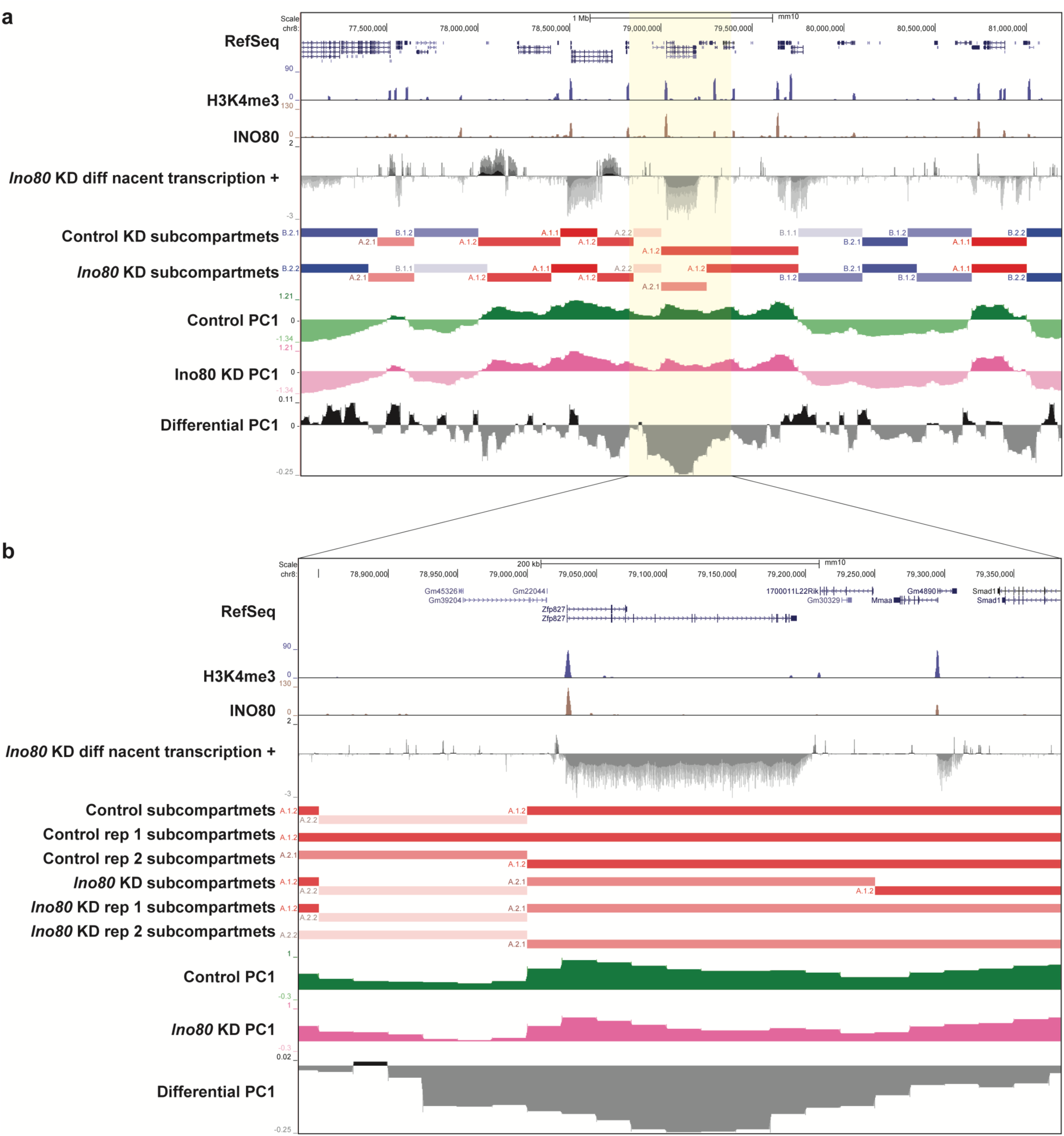
Subcompartment and compartment changes over *Zfp827* in *Ino80* KD relative to control. **a)** UCSC genome browser track for 4 Mb on chromosome 8 depicting H3K4me3 ChIP-seq, (Chronis et al., 2017), INO80 ChIP-seq (Xue et al., 2017), *Ino80* KD relative to control TT-seq (Patty et al., 2025), control and *Ino80* KD subcompartment Hi-C data, and control, *Ino80* KD and differential compartment Hi-C data. **b)** Zoom in of *Zfp827* locus with data as in A and independent replicates for subcompartments shown.

**Figure S6.**
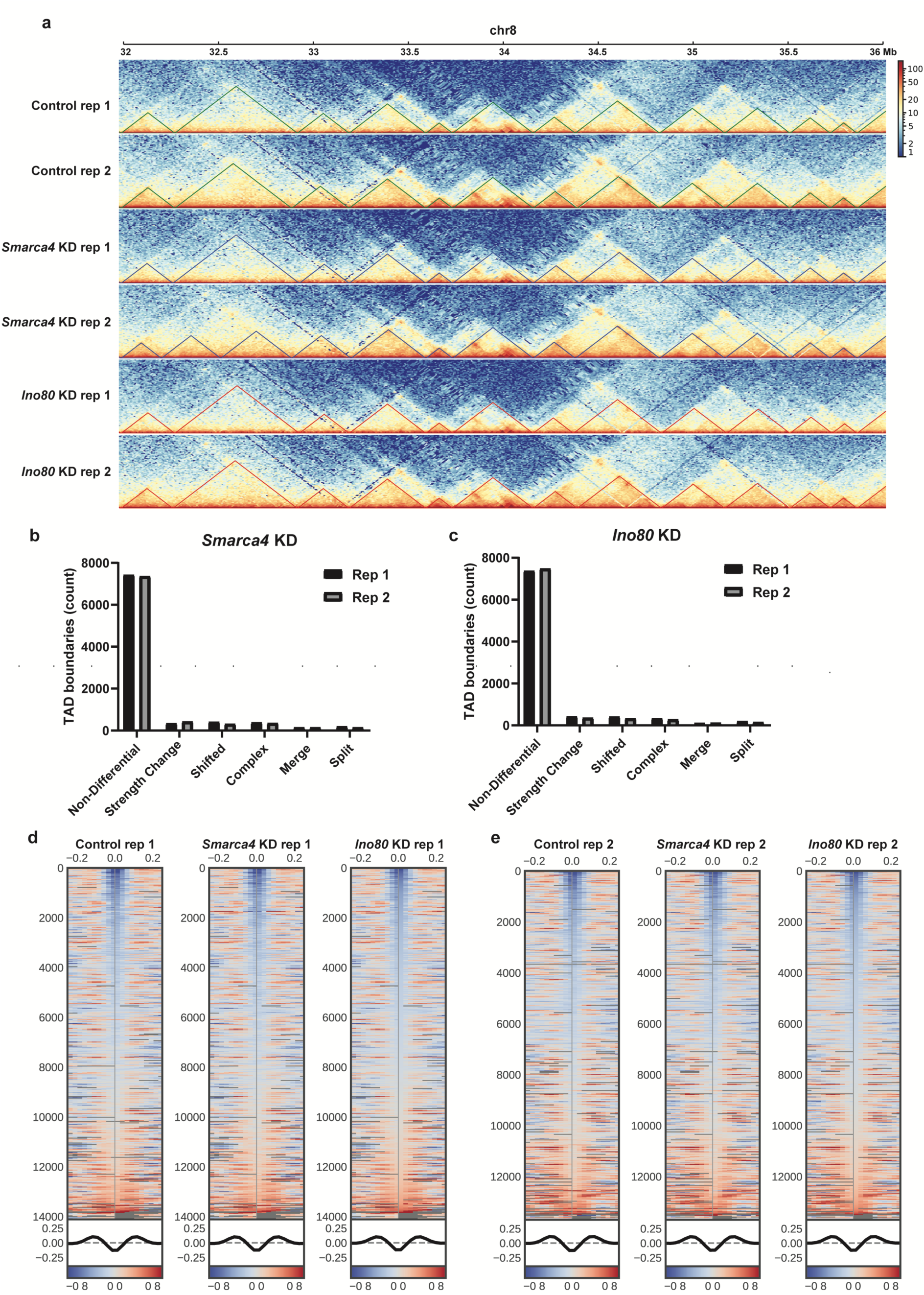
TAD interactions are reproducible across biological replicates. **a)** TAD interaction frequencies generated using HiCExplorer at 10 kb resolution over a 4 Mb portion of chromosome 8 for control (top) *Smarca4* KD (middle) and *Ino80* KD (bottom) individual replicates. **b-c)** Differential TAD boundary assignments in replicates for *Smarca4* KD (b) or *Ino80* KD (c) relative to control. **d-e)** Heatmaps depicting insulation score over all TAD boundaries in replicate 1 (d) and replicate 2 (e) for control (left) *Smarca4* KD (middle) and *Ino80* KD (right) generated using cooltools.

**Figure S7.**
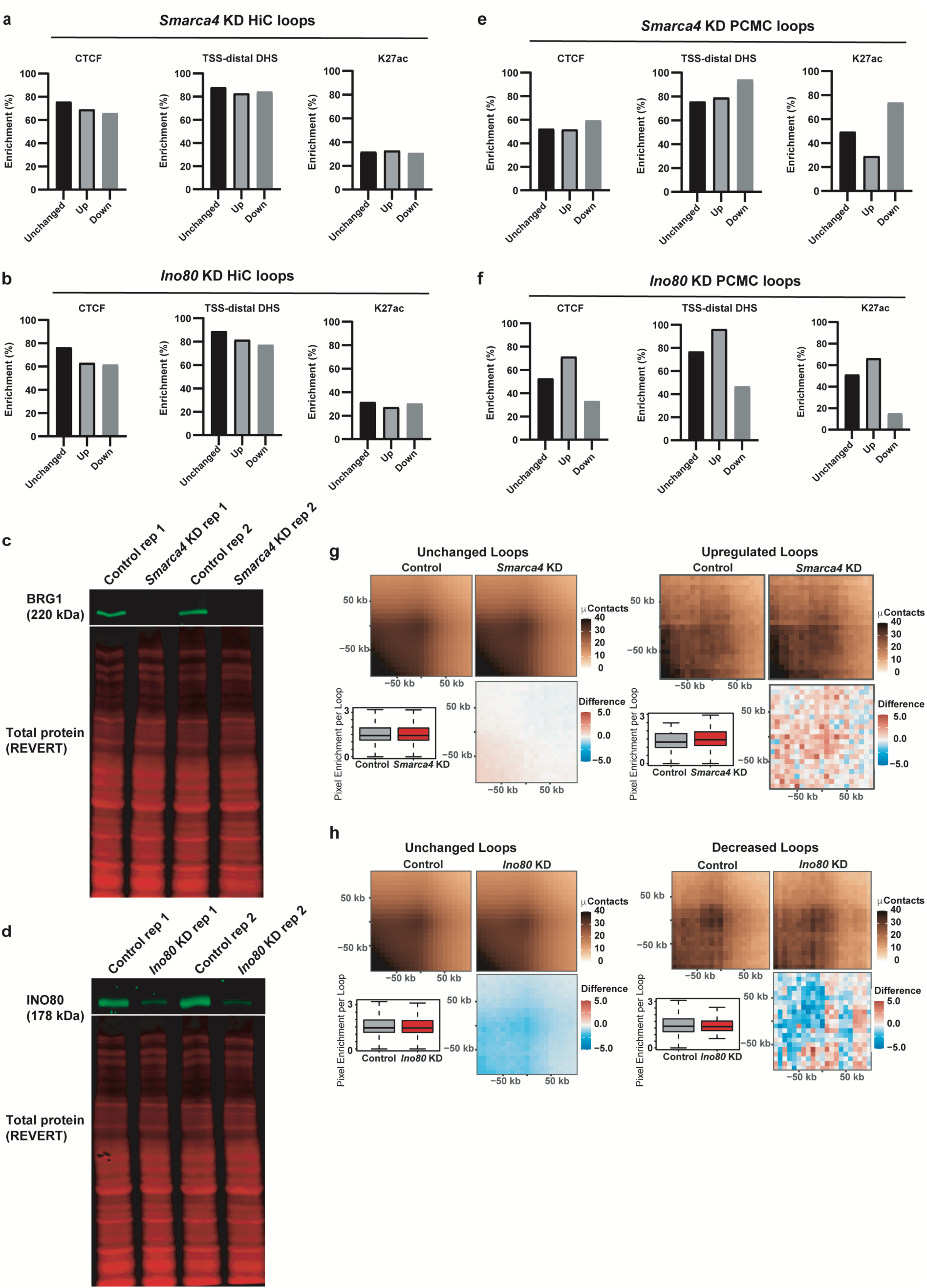
Differential PCMC loops are enriched for active enhancer marks. **a)** Percentage of unchanged, increased, or decreased loops in *Smarca4* KD Hi-C loops relative to control bound by (from ChIP-seq, left) (Ho et al., 2009), TSS-distal DHSs (from DNase-seq; middle) (ENCODE Project Consortium, 2012), and H3K27ac (from ChIP-seq, right) (Chronis et al., 2017). **b)** As in a, for *Ino80* KD Hi-C loops. **c-d)** Western blot analysis of BRG1 (c) and INO80 (d) upon esiRNA-mediated depletion of *Smarca4* or *Ino80*, respectively. REVERT total protein is used as a loading control. **e)** As in A, for *Smarca4* KD PCMC loops. **f)** As in a, for *Ino80* KD PCMC loops. **g)** Aggregate peak analysis (APA) for unchanged (left) and increased (right) loops in *Smarca4* KD (top right) relative to control (top left). Quantified peak enrichment (bottom left) and differential interaction frequency signal (bottom right) are also shown. **h)** As in e for unchanged (left) and decreased (right) loops in *Ino80* KD.

**Table S1.**
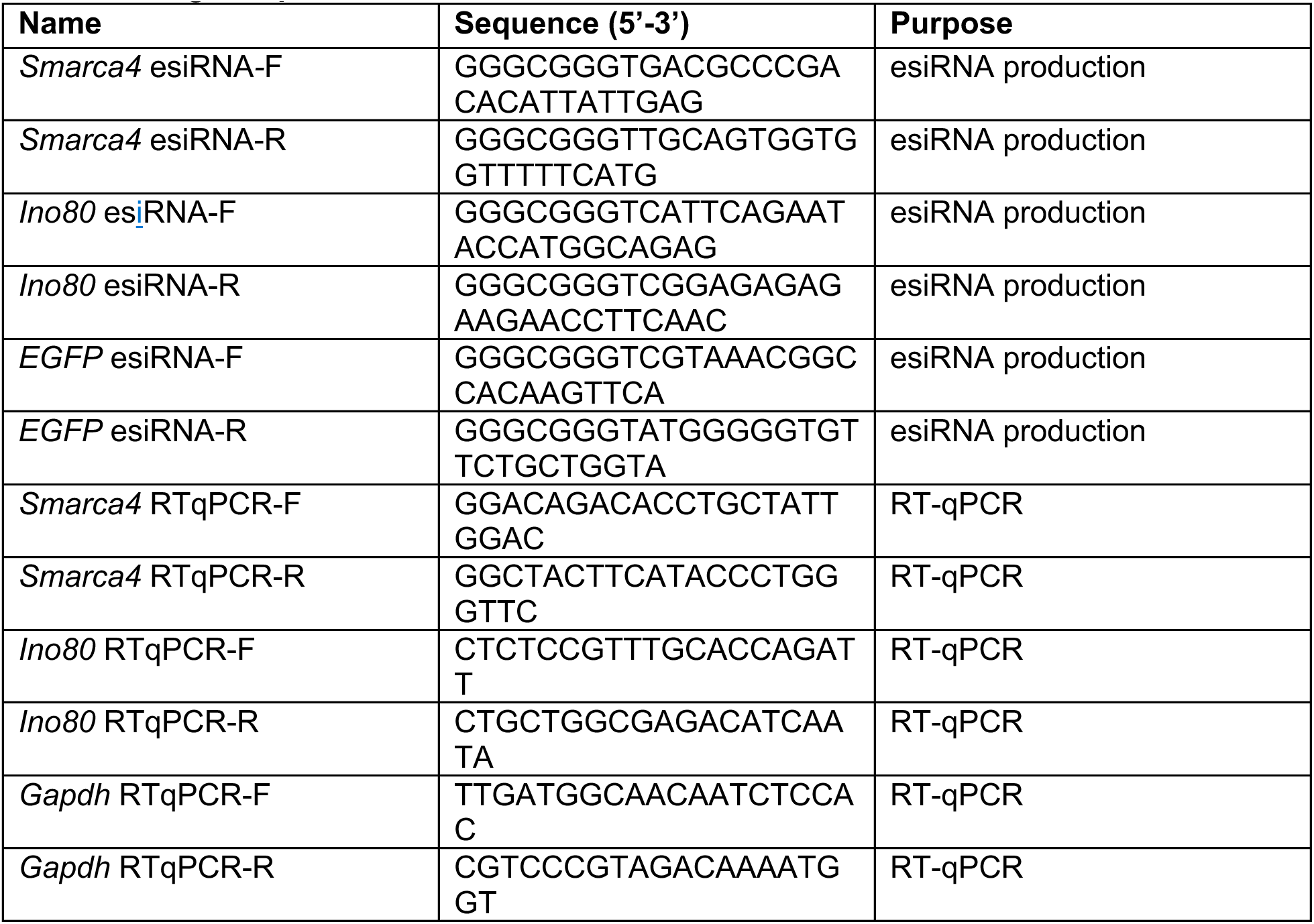
Oligo sequences.

**Table S2.** Publicly available ChIP-seq and CUT&RUN datasets.

| Dataset | SRA/GEO | Reference |
| --- | --- | --- |
| H3K4me1 | SRR5077636 | (Chronis et al., 2017) |
| H3K4me3 | SRR5077628 | (Chronis et al., 2017) |
| H3K27ac | SRR5077644 | (Chronis et al., 2017) |
| CTCF | SRR001985,<br>SRR001986,<br>SRR001987 | (Chen et al., 2008) |
| BRG1 | GSM358237,<br>GSM358238,<br>GSM358240 | (Ho et al., 2009) |
| INO80 | SRR5309424 | (Xue et al., 2017) |
| OCT4 | SRR7965412 | (Hainer et al., 2019) |
| NANOG | SRR6782491 | (Hainer et al., 2019) |
| SOX2 | SRR7965414 | (Hainer et al., 2019) |
| SUZ12 | SRR7965426 | (Hainer et al., 2019) |

